# vFLIM: Machine Learning-enabled Light Sheet Fluorescence Lifetime Imaging

**DOI:** 10.64898/2026.08.25.747039

**Authors:** Chad M. Hobson, Owen F. Puls, Jesse S. Aaron, Nicolas Denans, Anja Schmidt, Helen Farrants, Eric R. Schreiter, Teng-Leong Chew

**Author notes:** These authors contributed equally (Chad M. Hobson, Owen F. Puls).

## Abstract

The lifetime of fluorescent molecules provides an orthogonal readout to fluorescence intensity, opening experimental possibilities of measuring changes in local molecular environments, mechanical tension, and metabolism, among other factors. These changes are best studied live and *in vivo*; however, limitations of slow imaging speeds, high phototoxicity, and increased data size and complexity have significantly impeded progress on this front. Here, we present a complete and transferable pipeline consisting of a light sheet FLIM microscope and an accompanying machine learning model for data processing that renders long-term and/or high-speed volumetric FLIM (vFLIM) tractable in living systems. We benchmark this pipeline across several biological use cases, model systems, lifetime ranges, and spatiotemporal scales, showcasing a suite of possibilities that our workflow enables. This comprehensive pipeline from imaging to analysis is a crucial step forward towards disseminating the power of live vFLIM to the broader bioimaging community.

## Introduction

Owing to its high-contrast, molecule-specific data, rich in spatial and temporal context, fluorescence microscopy is a fundamental technique in the modern biologist’s toolkit^1–3^. Relying on intensity as the sole experimental readout, however, does not take full advantage of the wealth of information encoded in the fluorescence emission process. Recent years have seen a growth of techniques leveraging alternative properties of fluorophores. Of particular interest is fluorescence lifetime imaging microscopy (FLIM), which measures the characteristic relaxation time of a fluorophore from its excited state during the emission process^4,5^. This orthogonal readout has been broadly leveraged for applications including improved fluorescence multiplexing^6–9^, measuring local changes in microenvironments^10–13^, metabolic sensing^14–17^, voltage imaging^18–21^, and molecular tension measurements^22–25^. The extension of FLIM to time-series, volumetric acquisitions in live samples, however, has been hampered by several practical limitations.

Most notably, determining the lifetime of a fluorescent molecule necessitates measurements that encode temporal information. That is, one must measure the arrival of photons at different points in time relative to a known reference point – often a short pulse of excitation light. This process is therefore inherently slower than conventional intensity imaging. The prolonged measurement time also translates to increased sample irradiation, which accelerates photobleaching and toxicity. Finally, FLIM systems have conventionally been implemented onto laser scanning confocal microscopes, which are already known for their slow speed and high light dose^3,26^.

Recent advances in detector technology, however, have eased these constraints. The advent of Single-Photon Avalanche Diode (SPAD) array detectors^27–29^, for example, merge the photon-counting sensitivity of avalanche photodetectors (APDs) with the parallelization of cameras. These SPAD detectors, among other tools^21,30–33^, have enabled the realization of widefield FLIM systems, which measure lifetimes simultaneously across the field of view as opposed to point-by-point. While this dramatically improves speed, it does not necessarily minimize the issues of toxicity and photobleaching. Several groups have therefore coupled widefield FLIM detection and light sheet excitation^34–44^, exploring variations in (i) light sheet generation, (ii) detector configuration, (iii) lifetime measurement approaches, and (iv) objective lens configurations. By preserving image contrast while reducing overall photon dose, these demonstrations highlight the feasibility of time-lapse volumetric FLIM (vFLIM) of living samples. However, adoption of these systems as routine instruments has been comparatively limited.

The usability of these systems is low, in part, due to the sheer scale and complexity of the resulting data. Light sheet imaging alone generates terabytes of data in single imaging sessions^45^ – the additional dimension of lifetime only exacerbates this bottleneck. Moreover, before the experimental data becomes useful, lifetimes must be computed from the raw images. This computationally intensive step poses a prohibitive barrier to practical implementation. Previous work has begun exploring machine-learning (ML) approaches to tackle the computational work of lifetime extraction^46–50^, but they have yet to be adapted to the scale of light sheet data sets.

Here, we present an end-to-end and transferrable time-lapse vFLIM pipeline, incorporating a light sheet FLIM microscope and ML-based lifetime processing. Our hardware design builds on previous literature^34–44^, featuring widefield detection through a SPAD array detector, two-color static light sheet excitation generated by a Powell lens, and capillary-based sample mounting. For lifetime processing and analysis, we similarly adapted a fit-free machine learning model^47^ for significantly accelerated lifetime estimation with improved accuracy. This makes the transition from a single, 2D image to 3D, time-lapse data tractable. We benchmark this pipeline across multiple (i) model systems, (ii) time scales, (iii) length scales, (iv) lifetimes, and (v) biological applications. Our work thus establishes a comprehensive workflow from imaging to analysis for live, vFLIM in complex systems.

## Results

### Machine Learning-enabled Light Sheet vFLIM Pipeline

Our pipeline for time-lapse vFLIM is comprised of two major components: a light sheet FLIM system and an ML lifetime processing software package. The light sheet FLIM system (Figure 1a,b) was built upon a Simultaneous Multiview (SiMView) microscope^51^ – a dual-sided excitation and emission light sheet microscope. However, the FLIM module presented here only uses one excitation arm and one detection arm. The full details and specifications of the system are available in the materials and methods. In brief, two pulsed lasers (488 nm and 560 nm, 5 – 80 MHz repetition rate) were coupled together through a dichroic mirror, expanded into a light sheet through a Powell lens, and integrated into the existing excitation light path through a polarizing beamsplitter. The addition of two separate laser lines gated by mechanical shutters enables multi-channel light sheet FLIM in a single experiment. A Powell lens light sheet generation approach was chosen to (i) improve uniformity over a cylindrical lens and (ii) minimize pile-up artifacts previously reported with digitally scanned light sheets^42^. Two slit apertures were used to control both the light sheet thickness as well as the field of view over which the light sheet excited the specimen, with the latter being particularly critical for tiled acquisitions. Two dual-axis galvanometer scanning systems controlled the tip, tilt, and translation of the light sheet at the specimen. This setup is necessary for assuring proper light sheet alignment for each channel at all depths throughout the specimen^52–54^.

**Figure 1:**
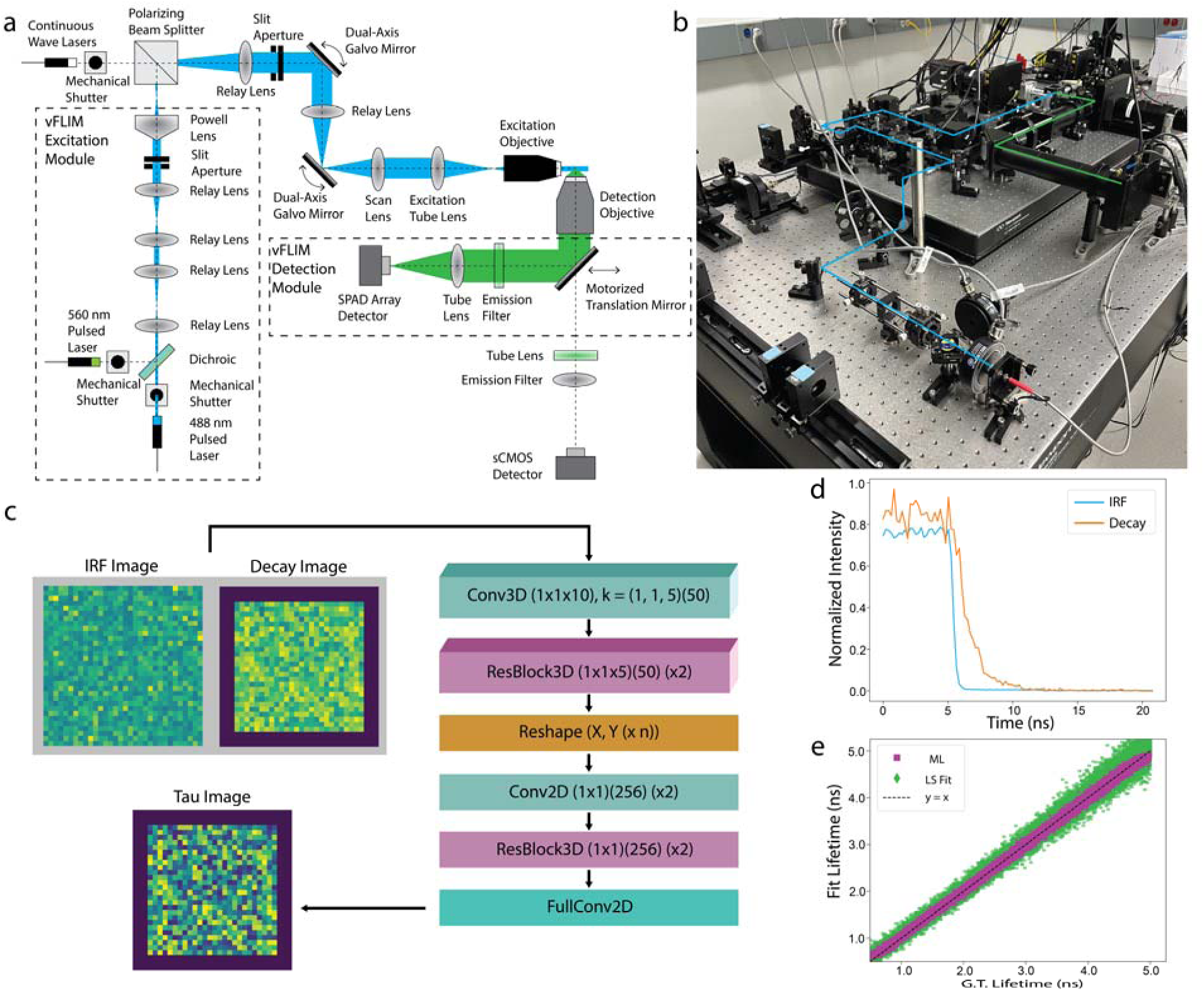
Light sheet FLIM system design and ML analysis pipeline. (a) Schematic overview of the light sheet FLIM excitation and detection paths. (b) Picture of the instrument with excitation (blue) and detection (green) paths overlaid. (c) Schematic of the ML pipeline. The architecture shown takes in the IRF and fluorescence decays as inputs, and outputs lifetime (tau) predictions. (d) Example traces of an IRF and decay taken from the same pixel in the inputs in (c). (e) ML and least-squares fitting predictions against ground-truth lifetimes for simulated test data.

For detection, we employ a 512×512 pixel SPAD array detector (Pi imaging). The pixel size of the SPAD array detector was matched to the original sCMOS detector used for intensity-based imaging. This allows for potential seamless integration of lifetime and intensity data sets without compromising field of view and efficiency of the SPAD array detector. In this work, however, we predominantly use the sCMOS detector for locating and positioning specimens as well as general system alignment. To switch between detection paths, we employ a motorized bypass mirror that is fully controlled by the instrument software. For measuring fluorescence lifetimes, we operate the detector in a “time-gated” imaging configuration (Figure S1). In brief, a detection “gate” is opened at a specific time delay from the laser pulse, during which each pixel measures the arrival of a photon or lack thereof. This process is repeated for a specified number of iterations, as prescribed by the “bit-depth”, to build up a greyscale image from individual binary images. The gate delay is then shifted, and the process repeated for a predetermined number of gate steps, thus sampling the characteristic timescale of fluorescence decay. Intensity images can be generated directly from the time-gated data by pixel-wise summing of the signal in each gate image, which is how intensity images in this work are constructed. In this configuration, it is critical to understand the pixel-wise system response for instantaneous decay (lifetime = 0), known as the “Instrument Response Function” (IRF), as the measured decay curve is the convolution of the IRF with actual underlying fluorescence decay. IRFs were experimentally measured for the appropriate wavelength prior to every experiment (Figure S1b), and spatial variation of the IRF (Figure S1c) was accounted for in any downstream processing and analysis.

Time-lapse vFLIM inherently poses a hurdle: data processing time. As described, this system produces a fluorescence decay curve at each of the ∼10^7 voxels for a 512×512×100 voxel volume. We reasoned that it would be ideal to calculate the lifetime at each voxel, without a priori knowledge, due to the theoretically intensity-independent nature of FLIM. However, initial attempts at processing even a single volume of this size required hours of computation with traditional curve-fitting methods (performed on a machine with specifications given in Methods). Extending this to time-series data sets would therefore yield lifetime predictions many days after the imaging experiment, significantly reducing the capacity for iteration and optimizations at the microscope.

To remedy this, the second pillar of our vFLIM pipeline adapts an existing machine learning (ML) framework^47^ to significantly improve processing efficiency and accuracy for modeling single-component decays (Figure 2a and Figure S2). The general framework for our ML pipeline is shown in Figure 1c. We made two fundamental changes to the original architecture^47^. First, we trained a model which considered the IRF as an additional input at every pixel. We reasoned that the model could learn the temporal relationship between the IRF and the response, leading to improved predictions. Indeed, including the IRF as a second channel increased the accuracy of the model by ∼20% (Figure 2b) without any significant additional computational burden. Second, to maximize imaging speed we reasoned that acquiring only the decay portion of the response—rather than the rise and the decay—would be sufficient for both measuring the fluorescence lifetime as well as producing intensity images of sufficient contrast. Figure 1d displays examples of such curves for the IRF and decay (compare to Figure 2c for examples of the complete system response). This increased the practical imaging speed by ∼2x, as well as decreased the size of the data by a similar factor. Interestingly, this also improved the accuracy of model predictions by ∼40% (Figure 2d). Moreover, while the training data was generated at a specific gate size (0.2094 ns), the model performed reasonably well over a range of gate sizes (Figure 2e-f and Figure S3). This advantage allows users to prioritize different aspects of FLIM experiments such as temporal sampling and speed without needing to retrain the prediction model.

**Figure 2:**
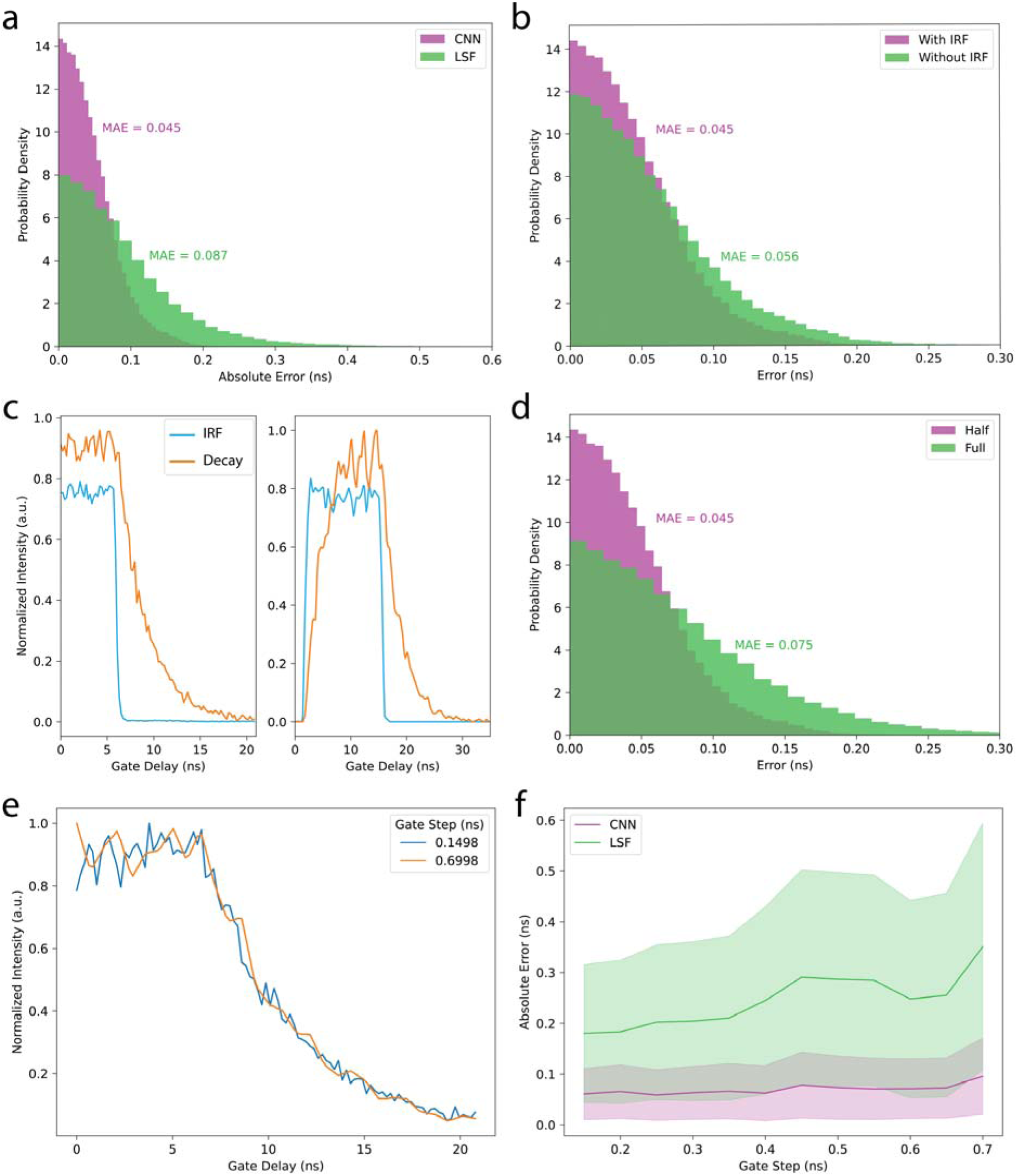
Advances in ML pipeline performance. (a) Comparison of lifetime prediction accuracy between the ML pipeline and least-squares fitting (LSF). The mean average error (MAE) is given for both. (b) Improvement in lifetime prediction accuracy by including the IRF as an additional input channel to the model. (c) Representative IRF and decays for the partial response (left) acquired and the full response of the system. (d) Improvement in lifetime prediction accuracy by passing the partial response as input to the model. (e) Representative decays simulated at two different gate step sizes. (f) Comparison of lifetime prediction accuracy for the ML pipeline and least-squares fitting over a range of gate step sizes. Note that the accuracy of the ML pipeline is reasonably stable over this experimentally relevant range.

Further, while fluorescence lifetime measurements often consider the photophysics of multi-component decays^5,55,56^, the mean lifetime—either an amplitude- or intensity-weighted average—is commonly reported^5,56^. This mean lifetime can be the result of many unique combinations of different amplitude-lifetime components because the parameters of multi-exponential decays are not truly independent for the purposes of curve-fitting^56^. This suggests that reporting an unambiguous single decay component is potentially more interpretable. Considering this, we chose to focus on a wide range of single-component decays (with lifetimes of 0.5-5.0ns) of varying noise levels, in contrast to more experiment-specific models^47^. This choice afforded flexibility in experimental design such that a wide array of fluorophores were equally available. Nevertheless, it is true that there will be some inherent error (∼0.0-0.5ns depending on the underlying decay) in using single-component models to predict underlying multi-component processes. However, for many of the purposes dominated by single-component decays displayed here, this is otherwise less critical.

In total, this ML pipeline drastically reduced the processing time for a single volume to minutes, not hours, facilitating a more rapid feedback loop between imaging and downstream results. For example, processing a ∼100GB dataset with curve-fitting on our local machine (see Methods) took ∼30hrs; the ML pipeline processed the same movie in ∼5hrs. Despite this already significant improvement, the underlying reality is even more stark due to two considerations. First, in both cases, a significant bottleneck remained loading the data. Second, the ML pipeline natively processes every pixel, while curve fitting methods require some amount of intensity threshold mask to ensure convergence. As a result, comparing the actual processing times highlights the significant efficiency increase: ∼300ms/slice for ML processing compared to ∼6s/slice for a masked slice (∼1% of pixels) via curve fitting. Moreover, the model demonstrated greater accuracy in predicting lifetimes than conventional least-squared curve fitting methods (both the prediction noise depicted in Figure 1e and the measured absolute error as shown in Figure 2a). We note that black-box models offer users less flexibility and tractability than explicit fitting functions. However, in this context, curve fitting is known to be especially sensitive to the gate delay at which the decay starts (Figure S4)^47,57^. As a result, this “flexibility” can adversely bias predictions; the parameter-free nature of the presented pipeline shields users from such artifacts. Taken together, the ML pipeline offers a powerful balance between robustness and processing speed that improves measured lifetime accuracy and overall makes light sheet FLIM experiments tractable.

### Lifetime-based Multiplexing

One of the common use cases for FLIM is improved multiplexing compared to conventional intensity imaging, as it enables spectrally overlapping fluorophores to be separated. To benchmark the performance of our pipeline in lifetime-based multiplexing, we conducted three experiments with steadily increasing levels of complexity.

First, we performed an *in vitro* control experiment by imaging two populations of green fluorescent beads with similar emission spectra, but with known, separable lifetimes and distinct diameters (∼2.1 ns and 2.0 µm; ∼4.0 ns and 4.19 µm) (Figure 3a-c, Movie S1)^58–61^. We could therefore separate the two populations of beads based purely on size to validate the effectiveness of our lifetime-based multiplexing. We observed clear lifetime separation that correlated with object size as expected (Figure 3a-c). Example intensity versus gate delay curves are shown in Figure 3c for the beads highlighted by white arrows in Figure 3a,b. The lifetimes of the beads across the populations (2.1 ± 0.1 and 4.0 ± 0.3 ns, respectively) (Figure 3c) are in good agreement with previously reported values^58–61^.

**Figure 3:**
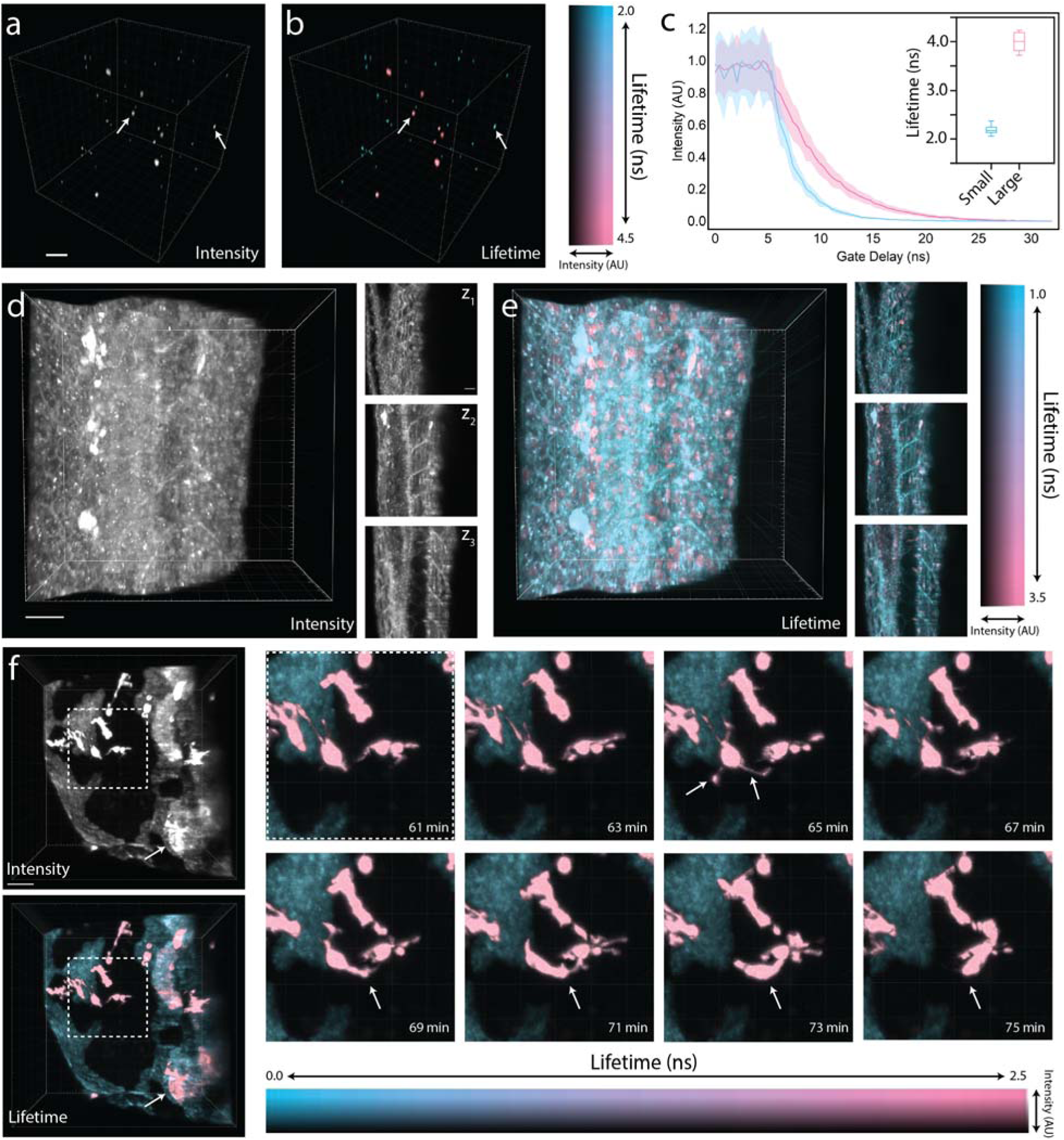
Lifetime-based multiplexing. 3D renderings of intensity (a) and intensity-modulated lifetime (b) for a light sheet vFLIM image of two populations of fluorescent beads with overlapping excitation spectra (scale bar = 30 µm). (c) Example decay curves for the beads highlighted by white arrows in (a) and (b). Shaded regions represent standard deviation of mean intensity. Inset shows quantification of lifetime for both populations of beads based on particle size. 3D renderings and individual z slices of intensity (d) and intensity-modulated lifetime (e) for a light sheet vFLIM image of trunk of zebrafish embryos fixed at 4 dpf expressing red fluorescent markers for cell membranes and nuclei (bactin2::mCherry-CAAX and bactin2::H2B-HaloTag labeled with Janelia Fluor 552, respectively) (volume rendering scale bar = 30 µm; z slice scale bar = 20 µm). (f) 3D renderings of intensity and intensity-modulated lifetime for a light sheet vFLIM time series of macrophages (mpeg1:EGFP) and tissue autofluorescence hindbrain of live zebrafish at 4dpf (scale bar = 30 µm). Dashed box is the region of interest for the timeseries shown on the right.

We then examined the trunk of zebrafish larvae fixed at 4 days post fertilization (dpf) expressing red fluorescent markers for cell membranes and nuclei (bactin2::mCherry-CAAX and bactin2::H2B-HaloTag labeled with Janelia Fluor 552^62^, respectively) (Figure 3d,e, Movie S2). A total image volume of ∼211 x 211 x 100 µm was acquired in approximately 20 s. In comparison to the *in vitro* fluorescent beads, this provides a specimen with broader lifetime variability and increased spatial overlap, with the addition of the optical artifacts associated with imaging deeper into biological tissue^63–65^. Despite these challenges, signals that are indistinguishable in intensity (Figure 3d) are easily separable by lifetime, as shown by intensity-modulated lifetime volume rendering and individual z slices (Figure 3e). A simple threshold at a lifetime of 2 ns is sufficient to split the single-channel stack into two separate volumes (Figure S5), effectively demonstrating the system’s capacity to multiplex based on lifetime in 3D biological tissue.

Given the gentle light sheet illumination scheme, the most applicable multiplexing use case for this system, however, is in *in vivo*, time-lapse, vFLIM. We imaged the dynamics of macrophages in the head of live zebrafish larvae at 4dpf (Figure 3f, Movie S3). Macrophages were labeled with a green fluorescent reporter (mpeg1:EGFP), which spectrally overlaps with the surrounding autofluorescence in the skin of the fish, complicating downstream segmentation and analysis. This is particularly relevant for immune cells residing at the surface of the brain, which are commonly obscured by autofluorescent signals (Figure 3f white arrow, Figure S6). While this multiplexing capacity is readily evident, we further extended its application to rapid and long-term volumetric imaging. Here, we demonstrate our capability to follow macrophages in motion at a rate of one 211 x 211 x 120 µm volume every 30 seconds (actual volume acquisition time of 9.2 seconds) for a duration of 3 hours (360 time points). We noted stable fluorescence intensity within macrophages despite considerable photobleaching of tissue autofluorescence. The specimen remained viable post-imaging, as evidenced by muscle twitching observed throughout the acquisition and continuous macrophage motility (Movie S3). The spatial resolution of the system (NA = 0.8) enabled detection of fine protrusions of the macrophage (Figure 3f inset, white arrow at t=65 minutes), yet with sufficient temporal resolution to support motion tracking (Figure 3f inset, white arrows from t=69 minutes to t=75 minutes). This demonstrates the ability of our integrated light sheet FLIM system and ML processing pipeline to perform high contrast, high resolution, vFLIM multiplexing for hundreds of time points at rates unattainable on conventional confocal systems.

### Multi-color, Time-lapse FLIM with Photoconversion

A current limitation of previous light sheet FLIM systems is their inability to excite and detect multiple wavelengths in the same experiment. This is, in part, due to complications in both hardware and software integration. Specifically, synchronization of excitation pulses, light sheet alignment, and differences in FLIM acquisition parameters between wavelengths make adding multiple spectral channels non-trivial. We solved the pulse synchronization issue by using the 488 nm laser as the master clock to provide an external trigger to both the 560 nm laser as well as to the SPAD array. Light sheet co-alignment across multiple wavelengths throughout the specimen depth is achieved by wavelength-dependent tuning of the galvanometer scanning mirror pair conjugated to the back focal plane of the excitation objective lens. The Powell lens used to generate the light sheet is not chromatically corrected, resulting in wavelength-dependent shifts in light sheet position. To compensate, the excitation objective lens position is adjusted via a piezo stage. Finally, we integrated software control to dynamically change FLIM acquisition parameters such as the gate offset, width, step size, and number as a function of wavelength on a per-stack basis. Together, these design features enable us to maximize image contrast while simultaneously optimizing the sampling of lifetime signatures across multiple spectral channels in one experiment.

As a demonstration, we performed time-lapse vFLIM on 4dpf live zebrafish larvae expressing a red fluorescent marker for blood vessels and a green-to-red photoconvertible marker for macrophages (fli1a:RFP-CAAX, mpeg1:Dendra2) (Figure 4, Movie S4). As in the previous experiment, we focused our attention on the head of the fish where tissue autofluorescence spectrally overlaps with the non-photoconverted macrophage population. A total of 61 time points were imaged every 2 minutes; at the halfway point in the timelapse, an external UV laser (375 nm) was used to broadly photoconvert the macrophages from only green to both green and red. The incomplete extinction of the green signal during photoconversion has been previously documented^66,67^. All 4 fluorescent structures (blood vessels, pre- and post-conversion macrophages, and autofluorescence) were clearly distinguishable via their lifetime signatures (Figure 4a), highlighting simultaneous lifetime multiplexing across spectral channels. We next used the lifetime information in the 488 nm channel to separate the macrophages from the tissue autofluorescence and segment them in 3D over time (Figure 4b, white dashed lines). The mask of the macrophages from the 488 nm channel was then applied to the 560 nm channel, and the average intensity was measured over time across both channels. A clear increase in intensity is visible in the 560 nm channel at the time of photoconversion (Figure 4c). This experiment highlights the feasibility of using targeted photoconversion to study behaviors of cellular subpopulations in relation to other spectrally-overlapping structures – an experiment not possible on conventional intensity-based systems or other light sheet FLIM microscopes.

**Figure 4:**
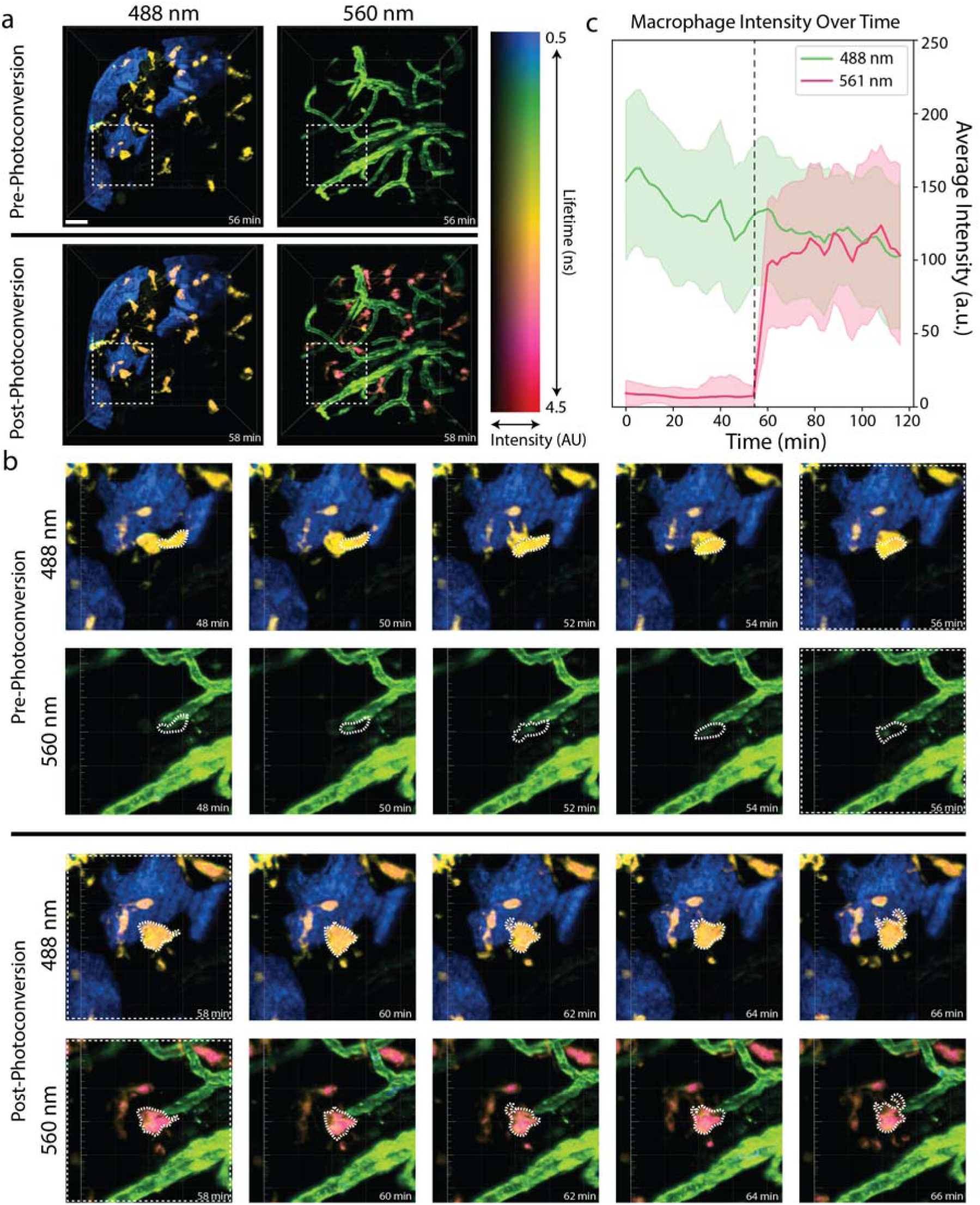
Multi-color, time-lapse FLIM with photoconversion. (a) 3D renderings of intensity-modulated lifetimes from a light sheet vFLIM time series of 4dpf live zebrafish embryos expressing a red fluorescent marker for blood vessels and a green-red photoconvertible marker for macrophages (fli1a:RFP-CAAX, mpeg1:Dendra2) (scale bar = 30 µm). Shown are the time points immediately pre- and post-photoconversion for both excitation channels (488 nm and 560 nm). (b) Time series of the inset shown in (a) for both excitation channels pre- and post-conversion. White dashed lines are representations of segmented macrophages based on the 488 nm channel for visualization purposes only. (c) Average and standard deviation of intensity of both excitation channels plotted over time for the segmented macrophages. The time of photoconversion is marked by the vertical dashed line.

### Large-volume, Long-term FLIM of Embryogenesis

A common application of light sheet microscopy is the capacity for gentle, long-term imaging of developing specimens over large fields of view. Potentially due to increased light dose compared to convention light sheet imaging and the small field of view of FLIM detectors, such experiments have yet to be demonstrated by any previous light sheet FLIM systems. To illustrate this unique capability, we imaged cell nuclei expressing a red fluorescent reporter (His2Av-mRFP) in a developing *Drosophila melanogaster* embryo (Figure 5, Movie S5). Volumetric images were collected every 4 minutes for 12 hours, spanning developmental stages of approximately 6 – 16^68,69^. This experiment would conventionally be limited by the limited field of view of current SPAD array detectors. To circumvent this, we integrated tiled FLIM acquisitions; in this instance, we collected 3 tiles along the anterior-posterior axis resulting in a total volume of approximately 211 x 525 x 202 µm, thus maintaining the full length of the embryo within the field of view over the entire imaging duration (Figure 5a). Tile alignment and stitching were performed based on fluorescence intensity and subsequently applied to the lifetime data. The light sheet was confined via a slit aperture to only excite the portion of the sample visible to the SPAD array detector, which was critical for avoiding excessive bleaching or light dose in the overlapping areas. This did, however, introduce spatially periodic intensity variations along the light sheet that were corrected for post-acquisition. To highlight the low photodamage of the light sheet configuration, we plotted average intensity within the nuclei over time (Figure 5b) – no photobleaching is discernable over the 12-hour time lapse. In addition, we observed canonical developmental events such as germband retraction and head involution^68,69^ (Figure 5c,d, Movie S5).

**Figure 5:**
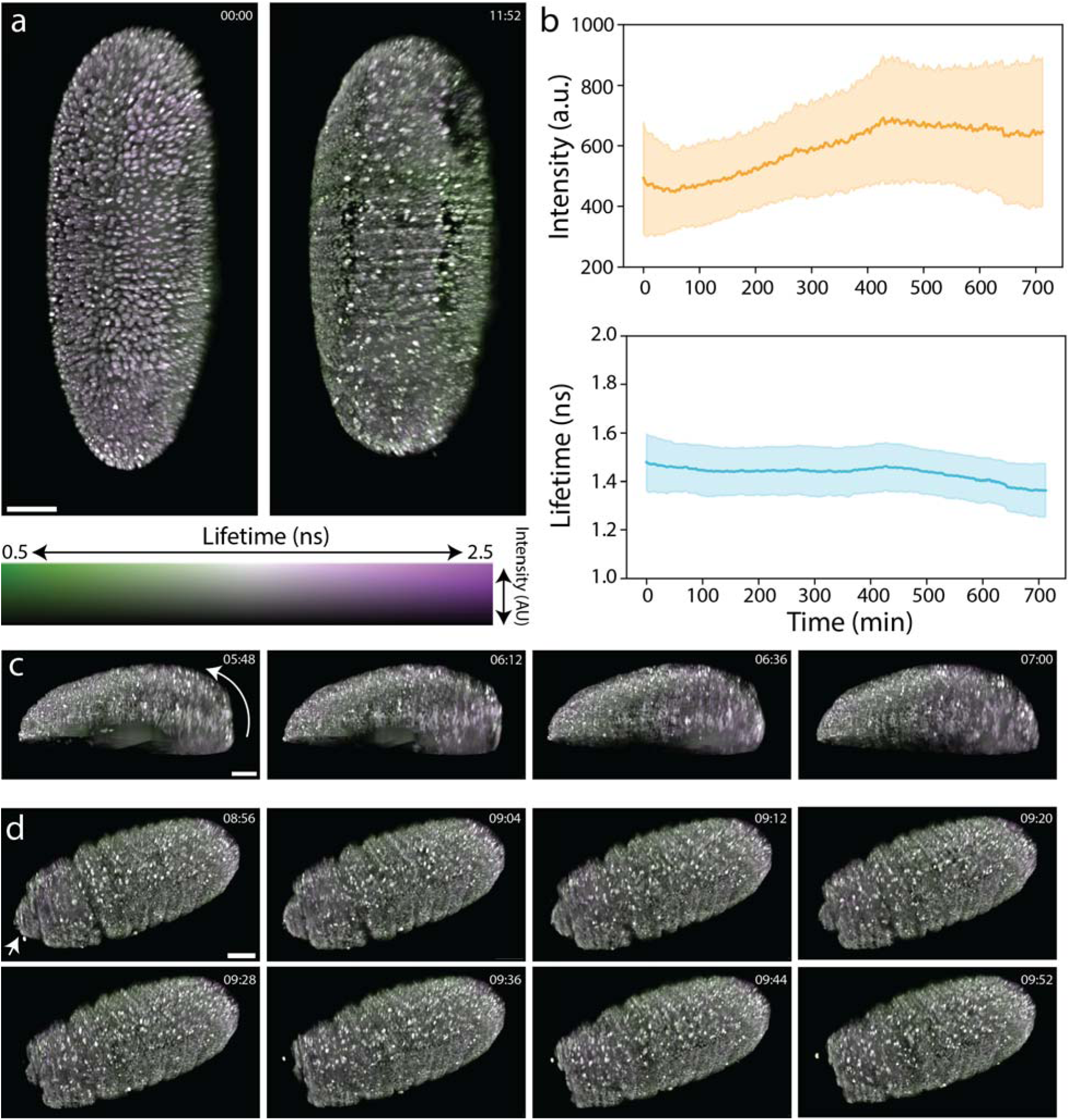
Tiled, long-term FLIM of embryogenesis. (a) 3D renderings of intensity-modulated lifetimes from a light sheet vFLIM time series of cell nuclei expressing a red fluorescent reporter (His2Av-mRFP) in a developing *Drosophila melanogaster* embryo. Shown are the ventral view of the first and final time points from the 12 hour imaging experiment over approximately developmental stages 6 - 16 (scale bar = 50 µm). (b) Average and standard deviation of intensity and lifetime of the nuclei plotted over time. (c) Time points during germband retraction (stages 12-13); ventral side of the embryo is at the top of the image (scale bar = 50 µm). (d) Time points during head involution (stages 14-16) shown from the ventral view (scale bar = 50 µm).

More than a demonstration of light sheet microscopy in the context of FLIM, this experiment provides three critical validations. First, we expect no significant changes in the lifetime of the fluorescent reporter as it is neither attached to a biosensor nor does the molecular environment of the nucleus dramatically change. Our results are in good agreement with this expectation as the lifetime over the entire imaging duration only fluctuates within the noise of the measurement (Figure 5b). Secondly, a hallmark claim of FLIM is that the lifetime measurement is independent of concentration^4^. If our system were to not uphold this claim, we would expect a striking correlation between changes in intensity and changes in lifetime. However, we see no such correlation throughout the time series (Figure 5b). Finally, we show that the measured lifetimes do not significantly vary with increased penetration depth relative to the light sheet, detection objective, or surface of the embryo (Figure S7), even in a highly scattering sample such as this. These three validations then provide significant confidence in the accuracy of our measured lifetimes and their independence of concentration, despite prolonged imaging during embryogenesis.

### High-speed FLIM of Calcium Transients

To benchmark our pipeline against the demands of fast, functional imaging, we performed vFLIM of spontaneous calcium biosensor (elavl3:NES-WHaloCaMP1a-EGFP-JF552) activity in the optic tectum of 4 dpf live zebrafish larvae (Figure 6). Each volume consisted of 5 z planes at a 4 µm spacing, imaged at a rate of one volume per 1.04 seconds. This proved fast enough to appropriately sample the pulsatile changes in lifetime due to calcium binding, while also maintaining the necessary sensitivity for detection. The previously reported mean lifetimes (amplitude-weighted) of WHaloCaMP1a-JF552 ([0.9, 2.9]ns for [EGTA, Ca^2+^]^70^) differ from those reported here by approximately ∼0.5ns, which is within the bounds of our expected error. Additionally, these previous measurements were made on purified protein with a 3-exponential fit, potentially explaining any further discrepancies. The volumetric nature of the experiment allowed us to track these transients at different depths in the optic tectum of the fish larvae, which is generally incompatible with conventional FLIM systems due to their slow acquisition speeds. This opens the door to rapid, *in vivo,* functional imaging using lifetime-based biosensors.

**Figure 6:**
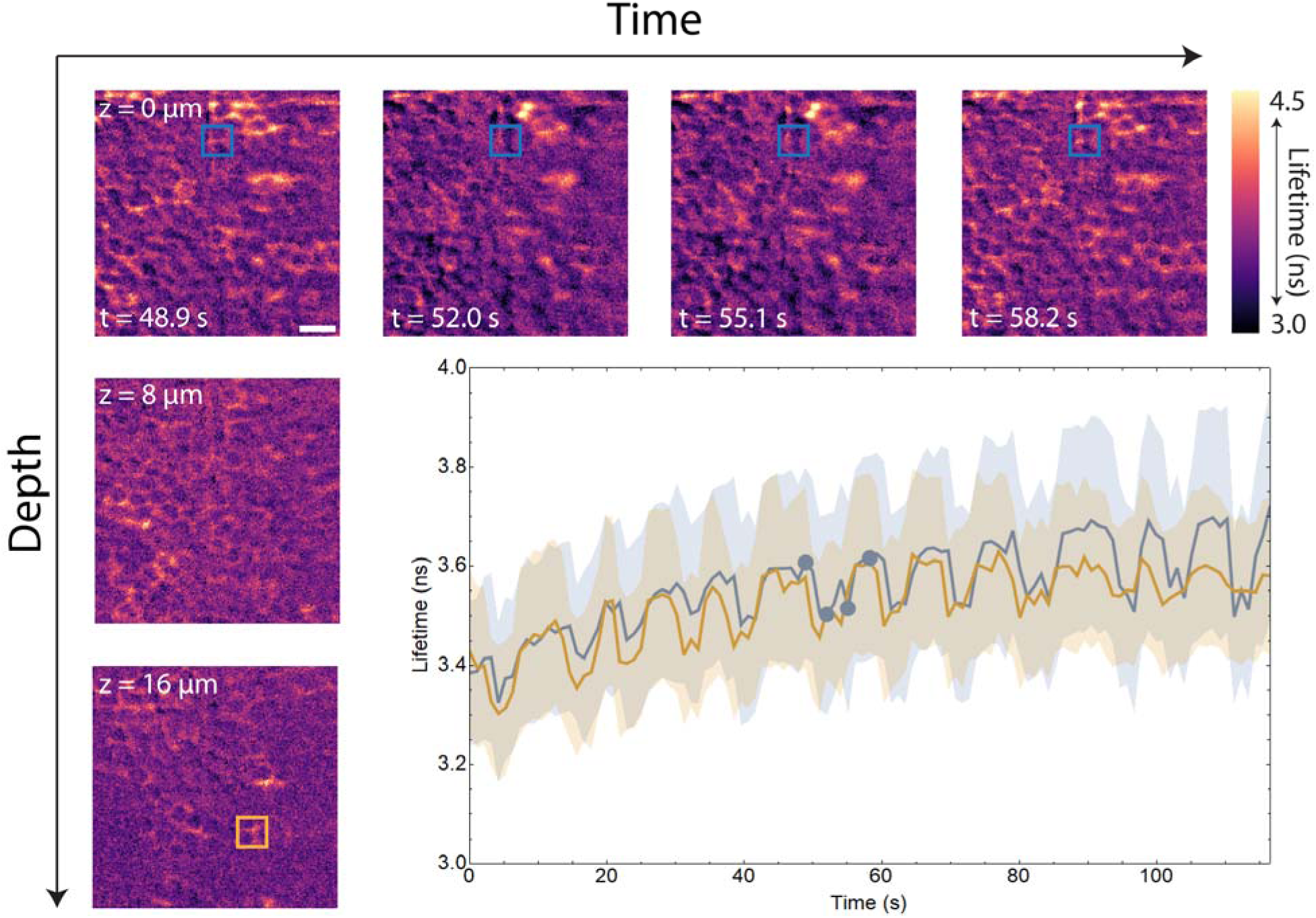
High-Speed FLIM of Calcium Transients. Volumetric, light sheet FLIM of a calcium biosensor (elavl3:NES-WHaloCaMP1a-EGFP) in the optic tectum of 4 dpf live zebrafish embryos. Shown are cropped fields of view (67 x 67 µm) for four time points for z = 0 µm, and three z planes at t = 48.9 seconds. Blue and orange boxes represent regions over which the lifetime versus time plot was created. Shown are average and standard deviation of lifetime. Blue dots on the graph correspond to the timepoint shown in the top row. Scale bar = 10 µm.

## Discussion

Fluorescence intensity measurement alone neglects the suite of orthogonal properties that fluorescent molecules encode. Doing so immediately restricts the scope of discovery before an experiment has even begun. This, however, encapsulates the current state of bioimaging. In particular, the lifetime of fluorescence molecules is often discarded or averaged out by conventional detectors, reducing the ability of the system to infer changes in biosensor activity or multiplex numerous sensors. FLIM circumvents this, but conventional systems are slow and phototoxic. Here, we present a light sheet adaption of FLIM to ameliorate these issues and bring lifetime imaging to living systems. The system reported here differentiates itself in several key areas from previous implementations^34–44^: (i) we employ a conventional SPIM geometry and mounting configuration^71^, which is convenient for imaging a variety of developing systems; (ii) the light sheet is generated via a Powell lens, which both increases uniformity compared to cylindrical lenses while minimizing the need for additional pile-up corrections associated with swept beams^42^; (iii) the hardware and software are designed to accommodate multi-channel FLIM excitation in the same experiment; and (iv) it features motorized switching between conventional light sheet and FLIM modes, permitting easier sample finding and the potential for multi-modal studies. Most importantly, our work extends beyond previous studies by benchmarking live imaging of multiple model organisms across length, time, and lifetime scales while highlighting varied applications of FLIM. Yet, failure to efficiently process the barrage of data will immediately cripple such applications.

Light sheet data present an imposing challenge due to sheer scale^45^. Multiplying this with the orthogonal lifetime readout therefore rapidly compounds the problem. First, accurate quantification of the fluorescence lifetime generally requires capturing sufficient photons while finely sampling the decay^57^, which is directly at odds with reducing imaging speed, data size, and phototoxicity. Post acquisition, fitting techniques often tradeoff between speed and accuracy, ranging from simple, fast, and less precise methods (rapid lifetime determination^72,73^), to mathematically complex but theoretically more precise techniques (convolutional methods^74,75^). Designing experiments to satisfy these constraints is non-trivial. Furthermore, processing this amount of data in a timely manner is equally arduous if the user is to iteratively test experimental conditions on the system.

To this end, we developed an ML pipeline in tandem with the hardware to mitigate the overall impact of these bottlenecks, offering a streamlined and efficient path from raw images to interpretable data. By leveraging GPU computations and the efficiency of feed-forward inference, this pipeline significantly reduces processing time, allowing users to quickly check imaging parameters, diagnose potential problems, and begin to interpret biological phenomena. Moreover, it removes the potential user bias in choosing fitting parameters. Because the training set includes decays at various levels of Poisson noise, the model predictions remain robust against low photon counts. Additionally, as evidenced throughout, the model produces accurate predictions at a range of gate delay steps. The combination of these two properties affords users flexibility in intelligently allocating the photon budget and the opportunity to minimize data size. Overall, this pipeline reduces many of the barriers to applying FLIM across biological scales.

Not only does our comprehensive pipeline constitute a concrete step forward in time-lapse, vFLIM, it also presents opportunities for future development. Lifetime imaging is, in general, still slow compared to conventional intensity-based systems. While we performed experiments reaching speeds of 60 ms per frame, traditional light sheet microscopes can image nearly an order of magnitude faster. While enabling, SPAD array detectors still suffer from comparatively low fill factors and small fields of view. Continued investment in developing detector technology will continue to address these limitations over time. For our implementation of light sheet FLIM specifically, future versions could take advantage of two-sided excitation and detection for imaging the full depth of scattering samples^51,76–78^, incorporate two-photon or UV laser lines for NADH and FAD lifetime imaging^14–17^, and improve the speed of motorized mode switching for rapid modality multiplexing.

Regarding the lifetime processing software, we trained the model on single-component exponential decays. While this assumption is sufficient to capture myriad phenomena, the photophysics of many fluorophores and sensors is more complex^5,6,10,11,34,55,70^. For most applications, however, a single lifetime offers a reasonable estimation of the mean lifetime, which is commonly the value reported in the literature^56^. Additionally, while we trained the model on a relatively large range of lifetimes (0.5-5.0 ns), this is not exhaustive. This problem, however, could be easily solved by fine-tuning the model with additional training data that exhibit longer lifetimes, where necessary.

Despite these challenges, the trajectory of lifetime imaging extends upward and the momentum it carries is palpable. The approach reported here demonstrates the possibility of *in vivo* experiments that can measure intramolecular tension or quantify tissue-specific metabolic changes without perturbing the very processes being studied. In turn, the possibility of such experiments allows biologists to think creatively about new hypotheses which were previously unfathomable. The existence of this future, however, hinges upon access and dissemination. The vFLIM pipeline we present here consists of a hardware design featuring readily available parts in a light sheet configuration common to both open-source and commercial products, and ML lifetime processing software that can be tuned, trained, and reapplied with minimal startup cost or investment in computing infrastructure. This and previous works therefore lay the blueprint for commercialized variants of light sheet vFLIM, which could become staples in labs and shared imaging facilities worldwide.

## Methods

### Instrument Design

#### Excitation Path

Two fiber-coupled pulsed lasers (50 mW, 5 – 80 MHz, <60 ps pulse width for VisUV-488 and <85 ps pulse width for VisUV-560, PicoQuant) are used for fluorescence excitation. The 488 nm line serves as the primary clock, providing a trigger signal to the 560 nm line to synchronize the pulses. Fiber outputs were collimated through a fiber coupler (CFC11A-A, Thorlabs) and gated by an individual mechanical shutter (LS6, Uniblitz) for each laser line. The couplers were rotated such that the polarization of the pulsed lasers were orthogonal to that of the continuous wave lasers in the existing SiMView microscope. Laser lines were combined via a dichroic mirror (DMSP505, Thorlabs) and twice expanded by a factor of 1.5x through a 50 mm (AC254-050-A, Thorlabs) and 75 mm (AC254-075-A, Thorlabs) as well as a 200 mm (AC254-200-A, Thorlabs) and 300 mm (AC254-300-A-ML, Thorlabs) lens pair, respectively. Light was relayed to an adjustable slit aperture (VA100, Thorlabs) to control the final thickness of the light sheet. A Powell lens (LOCP-8.9R05-3.0, Laserline Optics) was used to form a light sheet, after which the beam was coupled into the existing SiMView excitation path via a polarizing beam splitter (PBS251, Thorlabs). A 125 mm lens (49-361, Edmund Optics) was used to image the light sheet through a second variable slit aperture (SP 60, OWIS GmbH) which controlled the field of view over which the light sheet excited the specimen and onto a dual-axis galvanometer scanning system (6125LH, Cambridge Technology) conjugated to the sample plane to control tip/tilt of the light sheet. A second 125 mm lens (49-361, Edmund Optics) then re-imaged the sheet onto a second dual-axis galvanometer scanning system (6125LH, Cambridge Technology) conjugated to the sample plane to control the position of the light sheet at the sample. The sheet then passed through an 80 mm scan lens created by a pair of 150 mm lenses (47643, Edmund Optics) and through a final 125 mm excitation tube lens (49-361, Edmund Optics), and was imaged onto the specimen by a 6.4x water dipping excitation objective (54-12.5-31, Special Optics). The microscope control software (LabVIEW) and sample mounting hardware have been documented previously^51–53^.

#### Detection Path

Emitted fluorescence was collected through a 16x 0.8 NA water immersion detection objective (CFI75, Nikon). An elliptical mirror (BBE1-E02, Thorlabs) attached to a motorized stage (LGA201S06-B-TDBA-038, Nanotec) was used to redirect the fluorescence to the FLIM detection path. The position of the motor was set using an optical encoder (NTO4L-05-B12-HC, Nanotec) and a matched controller (C5-E, Nanotec). Control over these hardware modifications were integrated directly into the existing LabVIEW microscope control software. Emitted fluorescence was filtered (FF01-520/35-25, Semrock; FF02-617/73-25, Semrock; FF01-523/610-25, Semrock) based on the experiment at hand, and a 500 mm tube lens (ACT508-500-A-ML, Thorlabs) formed the image of the sample onto the SPAD array detector (SPAD512, Pi Imaging). The 488 nm pulsed laser line also served as the primary clock for the SPAD array detector gate synchronization. An external 5 volt signal was used to trigger the acquisition of individual frames. The native detector software (cSPAD version 2.04, Pi Imaging) was used for operating the detector in the “Gated Imaging” functionality, via a LabVIEW remote command interface. Default procedures for the noise calibration, dead pixel calibration, breakdown calibration, and master-slave offset calibrations were performed^79^. No gain calibration for the detector was used and instead all gain calibration values were set to zero. Images were streamed directly from the USB connection to binary images performed through the LabVIEW control software to maximize acquisition speed.

### Machine Learning Pipeline

#### Hardware

For benchmarking purposes, all computations were performed on a Linux (Ubuntu) system equipped with an Intel® Xeon® w7-3465X processor, 512 GB RAM, and an NVIDIA RTX A4000 GPU with 16GB of vRAM. Pre- and post-processing was also performed on this machine. The machine learning pipeline was also configured to run equivalently on the Janelia compute cluster which utilizes various CPU-GPU configurations. However, any processing times quoted reflect results from the single machine, not the compute cluster.

#### Architecture

The machine learning architecture used here largely matches that presented in the source^47^ and was defined using Tensorflow. However, a second residual block after the initial 3D convolution layer was added to match the symmetry of the residual blocks deeper in the network. The mean-squared error was similarly used for the loss term. The network was trained using the Adam optimizer. The learning rate varied from 1e-3 to 1e-2 using a triangular (period = 4*total number of steps) schedule of exponentially decaying amplitude (adapted from github.com/brianmanderson/Cyclic_Learning_Rate). The training data was shuffled at the end of each epoch. While the number of epochs was defined to be 200, the training stopped early after 35 epochs due to a stagnant validation loss, defined as no improvement in loss greater than 1e-5 over five consecutive epochs.

#### Training Data Generation

Training data was generated using the same principles as defined in the source^47^ but adapted from MATLAB into Python, with some additional changes. Training data consisted of [32,32,100,2] pixel images in which one channel corresponds to the IRF and the other corresponds to the decay. Values for the lifetime and intensity are chosen from uniform distributions and assigned to each pixel. Lifetime values were chosen from the range [0.5-5.0] ns and the intensity from the range [250,1500]. Along a 2-pixel-wide border of the image, the intensity is set to zero. Everything that follows was performed pixel-wise. An IRF trace is assigned by randomly choosing coordinates within a 256×512 pixel IRF image (we focused on one vertical half of the image due to symmetry). The IRF was padded with small random values to match the maximum gate delay acquired on the system—in all data here, this value is 0.6998*30 = 20.994 ns. When only the decay portion of the IRF was acquired, the IRFs were also backwards padded with a reflection of the decay. This was performed to mimic the actual response on the microscope, even though only a portion of it was acquired. The IRF was then normalized by dividing by the summed intensity.

The IRF was then convolved with a single-component exponential decay of unity amplitude and lifetime as defined previously. At this stage, the resulting trace is sampled at the temporal resolution of the IRF (gate step = 0.0186ns). The resulting decay was then multiplied by the defined intensity and Poisson noise was added accordingly. This trace and the IRF were both trimmed to remove the effect of the backwards padding on the IRF, resampled at a specified temporal resolution (default gate step = 0.2098 ns), and then linearly interpolated to the standard number of gates (nG = 100). Finally, the decay curve is rounded to integer values to match the datatype acquired on the microscope and then normalized by dividing by the maximum intensity. To avoid negative values introduced by the interpolation step, both the IRF and decay were clipped to 0.

After these images were created, both the decay and IRF channels were aligned in time so that each decay begins at approximately the same gate delay. Using the input IRF image, a reference point was selected by determining the minimum gate delay at which the IRF “decay” begins on the system. A Savitsky-Golay filter (scikit-image) of order 3 and subset size 51 gate steps was applied at pixel across the time dimension to broadly smooth the response. The gate delay corresponding to the minimum of the second derivative of this smoothed response was recorded as the “decay” start. The earliest decay start across the field of view was chosen as the reference point. Both the IRF and decay data were then shifted in time based on the number of gates between the decay start at each pixel and this reference point. To maintain the correct number of gates, they were then zero-padded a corresponding amount.

The two-channel image and the ground-truth lifetime were used as training pairs for the network. This training/validation dataset consisted of n = 80000/20000 such pairs.

### Sample Preparation

#### Fluorescent Beads

For Figure 3a-c, two populations of green fluorescent beads (4.19 µm, FSDG006, Bangs Laboratories; 2.0 µm, L4530-1ML, Sigma-Aldrich) were diluted into at a ratio of 1:5000 into 10 ml of agarose (2% in MilliQ water, A2576, Sigma-Aldrich). Molten agarose was aspirated into a glass capillary with an inner diameter of 1.5 mm (1472609, Hilgenberg). Once agarose was solidified, approximately 1 mm of agarose was extruded from the glass capillary by plugging one end of the capillary with dental wax.

#### Zebrafish Larvae

All zebrafish experiments were conducted in accordance with animal research guidelines from the National Institutes of Health and were approved by the Institutional Animal Care and Use Committee and the Institutional Biosafety Committee of Janelia Research Campus. Larvae were reared using standard protocols in 14:10 light–dark cycles at 28.5° C.

Tg(T2[bactin2::H2B-HaloTag]; myl7:GFP) and Tg(T2[bactin2::mCherry-CAAX]; myl7:GFP) are kind gifts from Dr. Phillip Keller. Tg(elavl3:NES-WHaloCaMP1a-EGFP)^70^; Tg(mpeg1:EGFP)gl22^80^, Tg(fli1a:RFP-CAAX)pt505^81^ and Tg(mpeg1:Dendra2)uwm12^82^ were obtained from the Zebrafish International Research Center (ZIRC).

For labeling with Janelia Fluor dyes, dye-ligands were delivered to 4 dpf zebrafish larvae by adding the dye-HaloTag ligand (final concentration of 4 µM from a 2-mM stock in DMSO) to the system water for 2 h. The larvae were then washed 4 times for 1 h with fresh system water before mounting for light sheet imaging.

To mount embryos for imaging, embryos were first anesthetized with αBungarotoxin (Invitrogen, B1601). Fish larvae were placed with a minimal amount of water on the hydrophobic plastic lid of a 35 mm dish; any excess water was removed from the fish. 50 µL of 1 mg/mL αBungarotoxin was added to the larvae for 2 min and was then removed and replaced with fresh system water. Larvae were then transferred into a 35 mm dish with system water and allowed to recover for approximately 20 min. Once anesthetized, one embryo was transferred at a time directly on the hydrophobic plastic lid of a 35 mm dish. Any excess water was removed and 400 uL of liquid agarose (2% in system water, A2576, Sigma-Aldrich) was added to the embryo. A glass capillary with an inner diameter of 1.5 mm (1472609, Hilgenberg) was attached to a 20-gauge needle (Nordson EFD, 7018178) by inserting the needle into the capillary and wrapping the base of the capillary with parafilm. The embryo was gently aspirated tail first into the glass capillary using a 3 ml syringe (Becton, Dickinson and Company, 309657), and the agarose was allowed to solidify for 10 minutes. Once solidified, dental wax was pushed into the capillary to extrude the agarose, allowing the head of the embryo to be exposed for imaging. Any additional agarose above the head was trimmed away.

#### Drosophila melanogaster embryos

For imaging of Drosophila embryos expressing a Histone-RFP1, the following fly line was used: w*; P{w[+mC]=His2Av-mRFP}III.1 (23650, Bloomington Drosophila Stock Center). Flies were kept at room temperature, and embryo collections were done on grape juice agar plates at 25° C for 2-4 h.

The mounting procedure for fly embryos was adapted from previous work^83^. Before imaging, embryos were dechorionated in 50% bleach for 1.5 min, collected in nets and washed with water. The embryos were then spread on a fresh grape juice agar plate for staging. Cellularizing embryos were selected under a fluorescence stereoscope and transferred to a water drop in a plastic dish. Air bubbles attached to the embryo were carefully removed with a 30-gauge needle (Becton, Dickinson and Company, 305128). The embryo was then covered with liquid ultra-low gelling agarose (2% in water, Sigma, A2576) and drawn with a pipet into a glass capillary with an inner diameter of 1.5 mm (1472609, Hilgenberg, Germany) by attaching the capillary to a 200 µl pipette tip with parafilm. The embryo was centered in the agarose column with a sewing needle. After polymerization, the part of the agarose column containing the embryo was pushed out of the with dental wax and mounted on the microscope.

### Image Acquisition

#### Instrument Response Function (IRF)

Instrument Response Functions (IRFs) were collected by performing time-gated acquisitions of a powered milk solution (MilliQ water mixed with M17200-500.0, Research Products International), which directly scattered pulsed laser light onto the SPAD array detector. 800 gate steps were collected at a gate step size of 18.6 ps and gate width of 20 ns. The 8 bit imaging mode was used, and IRFs were collected individually for each wavelength with the corresponding gate offset prior to all experiments shown here. IRF measurements were repeated 21 times, after which the average of all measurements was taken to suppress noise in the IRF measurement. No emission filter was used in the detection path during IRF measurements.

#### Experimental Data

All imaging parameters for experiments performed in this work are summarized in Table S1. For fluorescent bead experiments shown in Figure 3a-c, volumes were acquired by scanning the detection objective lens in conjunction with the light sheet while leaving the sample still. For all other imaging experiments, stacks were collected by moving the sample stage while leaving the detection objective and light sheet fixed. For 488 nm excitation experiments, a green bandpass filter was used in the emission path (FF01-520/35-25, Semrock); likewise for 560 nm excitation experiments, a red bandpass filter was used (FF02-617/73-25, Semrock). For two channel excitation experiments, a green-red multi-bandpass filter was used (FF01-523/610-25, Semrock).

### Image Processing

#### Pre-processing

The decay data were pre-processed at each pixel. A Savitzky-Golay filter of order 3 and subset set 5 was applied to lightly smooth the data. A mask was then applied to remove pixels of extremely low intensity (<5 total counts across the decay). The images were then normalized by dividing by the maximum intensity at each pixel. To account for the spatial patterns in the IRF, the data were shifted pixel-wise such that the start of the decay was approximately uniform across the field of view, as described in the Training Data generation section above. To match the size of the training data, both the IRF and decay responses were then linearly interpolated to 100 gates and broken into 256 32×32 (xy) patches.

#### Destriping

Two methods of destriping the summed intensity (never lifetime) images were used. For every dataset other than that presented in Figure 5, the python package pystripe (https://github.com/chunglabmit/pystripe) was minimally updated to function with the current version of python and numpy. This consisted of changing a small amount of numpy syntax. Otherwise, the code was used as-written with sigma = [64,128], level = 10, and all other parameters set to their default values.

For the dataset in Figure 5, the nature of the striping artifacts was distinct, due to the confined light sheet used for this tiled acquisition, as explained in the main text. This ultimately meant that the destriping methods utilized previously did not remove the artifacts as before. To remedy this, a pseuo-flatfield correction was applied. For each timepoint, the max projection (F) of the summed intensity projection (C)—this is a proxy for the flatfield image—was blurred with a Gaussian filter (σ=10). A separate copy (D) of the first z-slice of the summed intensity image—this slice contains no biological signal and is a substitute for the “dark” image—was heavily blurred using a Gaussian filter (σ=50). A third image (G) was then calculated using the flatfield expression: 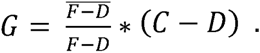 For this calculation, the quantity 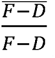 was clipped to a max value of 1000 to minimize artifacts. Finally, to minimize the prominence of regions outside the fly embryo, the corrected image (G) was processed at each z-slice in the following manner. Each z-slice of C was heavily blurred (σ=50) and an intensity threshold was defined from this blurred image using the scikit-image implementation of the Li method, resulting in a binary mask. This mask was processed using binary_fill_holes. Then the convex hull was calculated, dilated, and blurred slightly (σ=5), resulting in a pseudo-mask which consisted of pixels in the range [0,1]. G was then multiplied by this pseudo-mask. Finally, each timepoint was histogram-matched to the first timepoint to equilibrate the corrected intensities.

#### Stitching

The data in Figure 5 was stitched using the FIJI plugin BigStitcher^84^. The intensity channel alone was used to determine overlap and calculate fiducials. The stitching process proceeded as follows. First, tiles were roughly aligned using calculated pairwise shifts using the first timepoint. Interest points were then detected in every tile and timepoint using a difference-of-gaussian detector with sigma=2.7 and threshold=0.08. Using calculated correspondences, a two-step series of rigid and affine transformations were finally used to align adjacent tiles. These transformations were then duplicated onto the lifetime “channel”. Stitched volumes were fused using “Average Blending” and linear interpolation.

#### Intensity-Modulated Display

For visualization purposes, the summed intensity and lifetime were combined by creating an intensity-modulated display (IMD). First, the user selects a colormap. The colormap (in RGB format from matplotlib) was converted to HSV format, discretized into N=256 bins, and then duplicated and reshaped into a 256×256×3 array. This is a discretized representation of the colormap in HSV format, where each of the 3 channels corresponds to the hue, saturation, and value, accordingly. The chosen colormap will occupy the first (hue) channel. Similarly, a black-to-white colormap is discretized into N=256 bins, repeated and reshaped into a 256×256 array, and assigned to the last (value) channel of the HSV array. The saturation is kept identically at one. This HSV array is then converted back to RGB. This array is the colormap array.

After any destriping or other corrections to the intensity, the maximum display intensity was defined as a user-chosen percentage of the absolute maximum intensity (often >99%) over all time points, where relevant. The minimum display intensity could also be tuned but was kept at 0 throughout this work. Similarly, bounds on the minimum and maximum display lifetime were defined by the user. These two sets of bounds were then used to convert a two-channel, gray-scale image into an RGB image based on the chosen colormap. Two bin arrays of length N=256 were created with the lower- and upper-bounds defined by the intensity and lifetime bounds, respectively. These arrays correspond to equally spaced bins across each range. The lifetime and intensity images were then discretized using the corresponding bin arrays, resulting in arrays of indices indicating the position in the color and intensity ranges to which each pixel belong. An empty MxMx3 rgb image (where M is the x-y size of the input image) was then created and populated with rgb values drawn from the colormap array based on the bin indices.

#### Object Segmentation

For the data in Figure 4, macrophages were segmented using the FIJI plugin Labkit^85^. After predicting the fluorescence lifetimes, the entire timeseries from the 488 channel was converted to the BigStitcher-compatible file format HDF5 as a two-channel image: intensity and lifetime. This two-channel image was used to manually define masks for the macrophages in Labkit. These masks were subsequently used to train a pixel classifier using information from both channels (intensity and lifetime). After predicting the segmentation for the entire timeseries, the resulting binary images were used to mask both the 488 and 560 channels for macrophages. These masks were used to quantify the mean intensity over all macrophages in both channels over time, as shown in Figure 4c.

For the data in Figure 5, we calculated an intensity threshold for each timepoint of the stitched volume using the scikit-image implementation of local threshold method. This threshold was applied to roughly segment the nuclei. Objects of size unreasonable for nuclei (both large and small) were removed from the mask. This final mask was used to segment the nuclei. The mean lifetime across all nuclei was plotted in Figure 6b.

#### Curve Fitting

Decay starts were determined at each pixel as described in “Pre-processing” above. The collection of points along the curve after this point constitutes the “decay” for fitting purposes. Lifetimes were then calculated from the resulting decays in two separate ways. First, the scipy implementation of least-squares fitting was used to fit a single-component exponential decay of the form *F* = *Ae*^-*t*/*τ*^ + *C.* Predicted lifetimes and fit errors were reported. Error bounds on these predictions were calculated using the formula 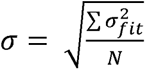 where N is the number of pixels in the region of interest. Second, the python library flimlib (https://github.com/flimlib/flimlib) was also used to determine the lifetime using an implementation of triple-integral curve fitting. The IRF was given as an argument to this fitting function, and the decay start was given as an estimate of the start of the fit.

#### Evaluating Model Performance

For evaluating model performance, additional test datasets of simulated decay images were generated using the same process as outlined above, with differences in parameter values noted were relevant. The absolute error of the predicted lifetime was calculated by taking the absolute value of the difference between prediction and ground truth lifetime.

## Data Availability

All raw image files underlying the images and movies shown here, as well as training and test datasets for the machine learning pipeline, are available in the following repository: 10.25378/janelia.33244515

## Code Availability

The machine learning model and additional software described here are available in the GitHub repository: https://github.com/aicjanelia/vFLIM.

## Acknowledgements

We thank: the current and former members of the HHMI Janelia Advanced Imaging Center team, Philipp Keller, and Hari Shroff for thoughtful input into this project; Johan Hummert (PicoQuant) and Harald Homulle (Pi imaging) for their assistance in hardware implementation; Andy Chiu (Sciotex) for his assistance in software implementation; Valentin Dunsing-Eichenauer and Pierre François Lenne (Aix-Marseille Université) for helpful discussions during the conception of this project; HHMI Janelia’s aquatics and invertebrate shared resources for assistance with zebrafish and Drosophila work; the Keller Lab (HHMI Janelia Research Campus) for sharing zebrafish lines; and Luke Lavis and the Janelia Open Science Initiative “open chemistry” team for providing the Janelia Fluor dyes used in this study.

## Author Contributions

C.M.H. and J.S.A. designed and constructed the light sheet FLIM system. O.F.P. developed the ML lifetime analysis software and performed all imaging processing and analysis. C.M.H., A.S., N.D., and H.F. prepared and performed imaging experiments. E.R.S. and T.-L.C. provided funding. E.R.S. and T.-L.C. oversaw experiments and the project. All authors contributed to writing and editing the manuscript.

## References

1. Cuny, A. P., Schlottmann, F. P., Ewald, J. C., Pelet, S. & Schmoller, K. M. Live cell microscopy: From image to insight. Biophysics Rev. 3, 021302 (2022).

2. Balasubramanian, H., Hobson, C. M., Chew, T.-L. & Aaron, J. S. Imagining the future of optical microscopy: everything, everywhere, all at once. Commun Biol 6, 1096 (2023).

3. Handbook Of Biological Confocal Microscopy. (Springer US, Boston, MA, 2006). doi:10.1007/978-0-387-45524-2.

4. Torrado, B., Pannunzio, B., Malacrida, L. & Digman, M. A. Fluorescence lifetime imaging microscopy. Nat Rev Methods Primers 4, 80 (2024).

5. Principles of Fluorescence Spectroscopy. (Springer US, Boston, MA, 2006). doi:10.1007/978-0-387-46312-4.

6. Bogdanova, Y. A. et al. Fluorescence lifetime multiplexing with fluorogen activating protein FAST variants. Commun Biol 7, 799 (2024).

7. Raymond, S. B., Boas, D. A., Bacskai, B. J. & Kumar, A. T. N. Lifetime-based tomographic multiplexing. J Biomed Opt 15, 046011 (2010).

8. Frei, M. S., Koch, B., Hiblot, J. & Johnsson, K. Live-Cell Fluorescence Lifetime Multiplexing Using Synthetic Fluorescent Probes. ACS Chem. Biol. 17, 1321–1327 (2022).

9. Frei, M. S. et al. Engineered HaloTag variants for fluorescence lifetime multiplexing. Nat Methods 19, 65–70 (2022).

10. Lakowicz, J. R., Szmacinski, H. & Johnson, M. L. Calcium imaging using fluorescence lifetimes and long-wavelength probes. J Fluoresc 2, 47–62 (1992).

11. Lakowicz, J. R., Szmacinski, H., Nowaczyk, K. & Johnson, M. L. Fluorescence lifetime imaging of calcium using Quin-2. Cell Calcium 13, 131–147 (1992).

12. Belashov, A. V. et al. Automatic segmentation of lysosomes and analysis of intracellular pH with Radachlorin photosensitizer and FLIM. Biochemical and Biophysical Research Communications 710, 149835 (2024).

13. Herrera-Ochoa, D., Pacheco-Liñán, P. J., Bravo, I. & Garzón-Ruiz, A. A Novel Quantum Dot-Based pH Probe for Long-Term Fluorescence Lifetime Imaging Microscopy Experiments in Living Cells. ACS Appl. Mater. Interfaces 14, 2578– 2586 (2022).

14. Skala, M. C. et al. In vivo multiphoton microscopy of NADH and FAD redox states, fluorescence lifetimes, and cellular morphology in precancerous epithelia. Proceedings of the National Academy of Sciences 104, 19494–19499 (2007).

15. Stringari, C. et al. Metabolic trajectory of cellular differentiation in small intestine by Phasor Fluorescence Lifetime Microscopy of NADH. Sci Rep 2, 568 (2012).

16. Winkler, U. & Hirrlinger, J. Crosstalk of Signaling and Metabolism Mediated by the NAD+/NADH Redox State in Brain Cells. Neurochem Res 40, 2394–2401 (2015).

17. Trinh, A. L. et al. Tracking Functional Tumor Cell Subpopulations of Malignant Glioma by Phasor Fluorescence Lifetime Imaging Microscopy of NADH. Cancers 9, 168 (2017).

18. Lazzari-Dean, J. R., Gest, A. M. M. & Miller, E. W. Measuring Absolute Membrane Potential Across Space and Time. Annual Review of Biophysics 50, 447–468 (2021).

19. Lazzari-Dean, J. R., Gest, A. M. & Miller, E. W. Optical estimation of absolute membrane potential using fluorescence lifetime imaging. eLife 8, e44522 (2019).

20. Gest, A. M. M., Grenier, V. & Miller, E. W. Optical Estimation of Membrane Potential Values Using Fluorescence Lifetime Imaging Microscopy and Hybrid Chemical-Genetic Voltage Indicators. Bioelectricity 6, 34–41 (2024).

21. Bowman, A. J., Huang, C., Schnitzer, M. J. & Kasevich, M. A. Wide-field fluorescence lifetime imaging of neuron spiking and subthreshold activity in vivo. Science 380, 1270–1275 (2023).

22. External strain on the plasma membrane is relayed to the endoplasmic reticulum by membrane contact sites and alters cellular energetics | Science Advances. https://www.science.org/doi/10.1126/sciadv.ads6132.

23. Pandzic, E., Whan, R. & Macmillan, A. Rapid FLIM Measurement of Membrane Tension Probe Flipper-TR. Methods Mol Biol 2402, 257–283 (2022).

24. Gonzalez-Garcia, M. C. et al. Exploring bacteria–surface interactions with a fluorescent membrane tension probe. Proceedings of the National Academy of Sciences 122, e2512977122 (2025).

25. Deo, C. Hybrid Fluorescent Probes for Imaging Membrane Tension Inside Living Cells. ACS Cent Sci 6, 1285–1287 (2020).

26. Icha, J., Weber, M., Waters, J. C. & Norden, C. Phototoxicity in live fluorescence microscopy, and how to avoid it. BioEssays 39, 1700003 (2017).

27. Zappa, F., Tisa, S., Tosi, A. & Cova, S. Principles and features of single-photon avalanche diode arrays. Sensors and Actuators A: Physical 140, 103–112 (2007).

28. Rochas, A. et al. First fully integrated 2-D array of single-photon detectors in standard CMOS technology. IEEE Photonics Technology Letters 15, 963–965 (2003).

29. Zappa, P. et al. Integrated array of avalanche photodiodes for single-photon counting. in 27th European Solid-State Device Research Conference 600–603 (1997). doi:10.1109/ESSDERC.1997.194500.

30. Bowman, A. J., Klopfer, B. B., Juffmann, T. & Kasevich, M. A. Electro-optic imaging enables efficient wide-field fluorescence lifetime microscopy. Nat Commun 10, 4561 (2019).

31. Hirvonen, L. M., Fisher-Levine, M., Suhling, K. & Nomerotski, A. Photon counting phosphorescence lifetime imaging with TimepixCam. Rev. Sci. Instrum. 88, 013104 (2017).

32. Lakowicz, J. R. & Berndt, K. W. Lifetime selective fluorescence imaging using an rf phase sensitive camera. Rev. Sci. Instrum. 62, 1727–1734 (1991).

33. Wang, X. F., Uchida, T., Coleman, D. M. & Minami, S. A Two-Dimensional Fluorescence Lifetime Imaging System Using a Gated Image Intensifier. Appl. Spectrosc., AS 45, 360–366 (1991).

34. Weber, P. et al. Monitoring of Apoptosis in 3D Cell Cultures by FRET and Light Sheet Fluorescence Microscopy. International Journal of Molecular Sciences 16, 5375–5385 (2015).

35. Mitchell, C. A. et al. Functional in vivo imaging using fluorescence lifetime light-sheet microscopy. Opt. Lett., OL 42, 1269–1272 (2017).

36. Funane, T. et al. Selective plane illumination microscopy (SPIM) with time-domain fluorescence lifetime imaging microscopy (FLIM) for volumetric measurement of cleared mouse brain samples. Rev. Sci. Instrum. 89, 053705 (2018).

37. Hirvonen, L. M. et al. Lightsheet fluorescence lifetime imaging microscopy with wide-field time-correlated single photon counting. J Biophotonics 13, e201960099 (2020).

38. Li, R. et al. Digital scanned laser light-sheet fluorescence lifetime microscopy with wide-field time-gated imaging. Journal of Microscopy 279, 69–76 (2020).

39. Nutt, K. J. et al. High-efficiency digitally scanned light-sheet fluorescence lifetime microscopy (DSLM-FLIM). 2023.06.02.543377 Preprint at 10.1101/2023.06.02.543377 (2023).

40. Samimi, K. et al. Light-sheet autofluorescence lifetime imaging with a single-photon avalanche diode array. J Biomed Opt 28, 066502 (2023).

41. Nedbal, J. et al. A time-correlated single photon counting SPAD array camera with a bespoke data-processing algorithm for lightsheet fluorescence lifetime imaging (FLIM) and FLIM videos. Sci Rep 14, 7247 (2024).

42. Dunsing-Eichenauer, V. et al. Fast volumetric fluorescence lifetime imaging of multicellular systems using single-objective light-sheet microscopy. Commun Biol 8, 1785 (2025).

43. Knight, V. R. et al. Fast wide-field light sheet electro-optic FLIM. *Opt. Express*, OE 34, 15065–15073 (2026).

44. Greger, K., Neetz, M. J., Reynaud, E. G. & Stelzer, E. H. K. Three-dimensional Fluorescence Lifetime Imaging with a Single Plane Illumination Microscope provides an improved Signal to Noise Ratio. *Opt. Express*, OE 19, 20743–20750 (2011).

45. Ouyang, W. & Zimmer, C. The imaging tsunami: Computational opportunities and challenges. Current Opinion in Systems Biology 4, 105–113 (2017).

46. Chen, Y.-I. et al. Generative adversarial network enables rapid and robust fluorescence lifetime image analysis in live cells. Commun Biol 5, 18 (2022).

47. Smith, J. T. et al. Fast fit-free analysis of fluorescence lifetime imaging via deep learning. Proceedings of the National Academy of Sciences 116, 24019–24030 (2019).

48. Zang, Z. et al. Fast Analysis of Time-Domain Fluorescence Lifetime Imaging via Extreme Learning Machine. Sensors 22, 3758 (2022).

49. Wang, Q. et al. Simple and Robust Deep Learning Approach for Fast Fluorescence Lifetime Imaging. Sensors 22, 7293 (2022).

50. Adhikari, M., Houhou, R., Hniopek, J. & Bocklitz, T. Review of Fluorescence Lifetime Imaging Microscopy (FLIM) Data Analysis Using Machine Learning. Journal of Experimental and Theoretical Analyses 1, 44–63 (2023).

51. Tomer, R., Khairy, K., Amat, F. & Keller, P. J. Quantitative high-speed imaging of entire developing embryos with simultaneous multiview light-sheet microscopy. Nat Methods 9, 755–763 (2012).

52. Royer, L. A. et al. Adaptive light-sheet microscopy for long-term, high-resolution imaging in living organisms. Nat Biotechnol 34, 1267–1278 (2016).

53. McDole, K., et al. In Toto Imaging and Reconstruction of Post-Implantation Mouse Development at the Single-Cell Level. Cell 175, 859–876.e33 (2018).

54. Hobson, C. M. et al. Practical considerations for quantitative light sheet fluorescence microscopy. Nat Methods 19, 1538–1549 (2022).

55. Fidler, V. & Kapusta, P. Fluorescence Kinetics and Time-Resolved Measurement. in Fluorescence Spectroscopy and Microscopy in Biology (eds Šachl, R. & Amaro, M.) 53–86 (Springer International Publishing, Cham, 2023). doi:10.1007/4243_2022_31.

56. Li, Y. et al. Investigations on Average Fluorescence Lifetimes for Visualizing Multi-Exponential Decays. Front. Phys. 8, (2020).

57. Fuhrmann, N., Brübach, J. & Dreizler, A. On the mono-exponential fitting of phosphorescence decays. Appl. Phys. B 116, 359–369 (2014).

58. Orthaus-Mueller, S., et al. rapidFLIM: The New and Innovative Method for Ultra fast FLIM Imaging.

59. Patting, M. et al. Fluorescence decay data analysis correcting for detector pulse pile-up at very high count rates. OE 57, 031305 (2018).

60. Houston, J. P., Naivar, M. A., Jenkins, P. & Freyer, J. P. Capture of Fluorescence Decay Times by Flow Cytometry. Current Protocols in Cytometry 59, 1.25.1–1.25.21 (2012).

61. Kapitany, V., Zickus, V., Fatima, A., Carles, G. & Faccio, D. Single-shot time-folded fluorescence lifetime imaging. Proceedings of the National Academy of Sciences 120, e2214617120 (2023).

62. Zheng, Q. et al. Rational Design of Fluorogenic and Spontaneously Blinking Labels for Super-Resolution Imaging. ACS Cent. Sci. 5, 1602–1613 (2019).

63. Reiche, M. A. et al. When light meets biology – how the specimen affects quantitative microscopy. J Cell Sci 135, jcs259656 (2022).

64. White, N. S., Errington, R. J., Fricker, M. D. & Wood, J. L. Multidimensional Fluorescence Microscopy: Optical Distortions in Quantitative Imaging of Biological Specimens. in Fluorescence Microscopy and Fluorescent Probes (ed. Slavík, J.) 47–56 (Springer US, Boston, MA, 1996). doi:10.1007/978-1-4899-1866-6_4.

65. Schwertner, M., Booth, M. J. & Wilson, T. Specimen-induced distortions in light microscopy. Journal of Microscopy 228, 97–102 (2007).

66. Gurskaya, N. G. et al. Engineering of a monomeric green-to-red photoactivatable fluorescent protein induced by blue light. Nat Biotechnol 24, 461–465 (2006).

67. Chudakov, D. M., Lukyanov, S. & Lukyanov, K. A. Tracking intracellular protein movements using photoswitchable fluorescent proteins PS-CFP2 and Dendra2. Nat Protoc 2, 2024–2032 (2007).

68. Atlas of Drosophila Development by Volker Hartenstein. https://www.sdbonline.org/sites/fly/atlas/00atlas.htm.

69. Bate, Michael. & Martinez Arias, A. The Development of Drosophila Melanogaster. *The Development of Drosophila melanogaster* (Cold Spring Harbor Laboratory Press, Plainview, N.Y, 1993).

70. Farrants, H. et al. A modular chemigenetic calcium indicator for multiplexed in vivo functional imaging. Nat Methods 21, 1916–1925 (2024).

71. Huisken, J., Swoger, J., Del Bene, F., Wittbrodt, J. & Stelzer, E. H. K. Optical Sectioning Deep Inside Live Embryos by Selective Plane Illumination Microscopy. Science 305, 1007–1009 (2004).

72. Ballew, R. M. & Demas, J. N. An error analysis of the rapid lifetime determination method for the evaluation of single exponential decays. Anal. Chem. 61, 30–33 (2002).

73. Woods, R. J., Scypinski, Stephen. & Love, L. J. Cline. Transient digitizer for the determination of microsecond luminescence lifetimes. Anal. Chem. 56, 1395–1400 (2002).

74. de Jong, F., Martín, C., Hofkens, J. & Van der Auweraer, M. Data Analysis Methods in Time-Resolved Fluorescence Spectroscopy: A Tutorial Review. Chemistry – A European Journal 31, e202401799 (2025).

75. O’Connor, D. V., Ware, W. R. & Andre, J. C. Deconvolution of fluorescence decay curves. A critical comparison of techniques. J. Phys. Chem. 83, 1333–1343 (1979).

76. Krzic, U., Gunther, S., Saunders, T. E., Streichan, S. J. & Hufnagel, L. Multiview light-sheet microscope for rapid in toto imaging. Nat Methods 9, 730–733 (2012).

77. Kumar, A. et al. Dual-view plane illumination microscopy for rapid and spatially isotropic imaging. Nat Protoc 9, 2555–2573 (2014).

78. Huisken, J. & Stainier, D. Y. R. Even fluorescence excitation by multidirectional selective plane illumination microscopy (mSPIM). *Opt. Lett.*, OL 32, 2608–2610 (2007).

79. cSPAD system manual | Pi Imaging. https://piimaging.com/doc-cspad.

80. Ellett, F., Pase, L., Hayman, J. W., Andrianopoulos, A. & Lieschke, G. J. mpeg1 promoter transgenes direct macrophage-lineage expression in zebrafish. Blood 117, e49–56 (2011).

81. Corti, P. et al. Interaction between alk1 and blood flow in the development of arteriovenous malformations. Development 138, 1573–1582 (2011).

82. Harvie, E. A., Green, J. M., Neely, M. N. & Huttenlocher, A. Innate immune response to Streptococcus iniae infection in zebrafish larvae. Infect Immun 81, 110–121 (2013).

83. Royer, L. A., Lemon, W. C., Chhetri, R. K. & Keller, P. J. A practical guide to adaptive light-sheet microscopy. Nat Protoc 13, 2462–2500 (2018).

84. Hörl, D. et al. BigStitcher: reconstructing high-resolution image datasets of cleared and expanded samples. Nat Methods 16, 870–874 (2019).

85. Frontiers | LABKIT: Labeling and Segmentation Toolkit for Big Image Data. https://www.frontiersin.org/journals/computer-science/articles/10.3389/fcomp.2022.777728/full.

