## Supplemental Materials for "vFLIM: Machine Learning-enabled Light Sheet Fluorescence Lifetime Imaging"

[†]These authors contributed equally

#### Supplemental Figures

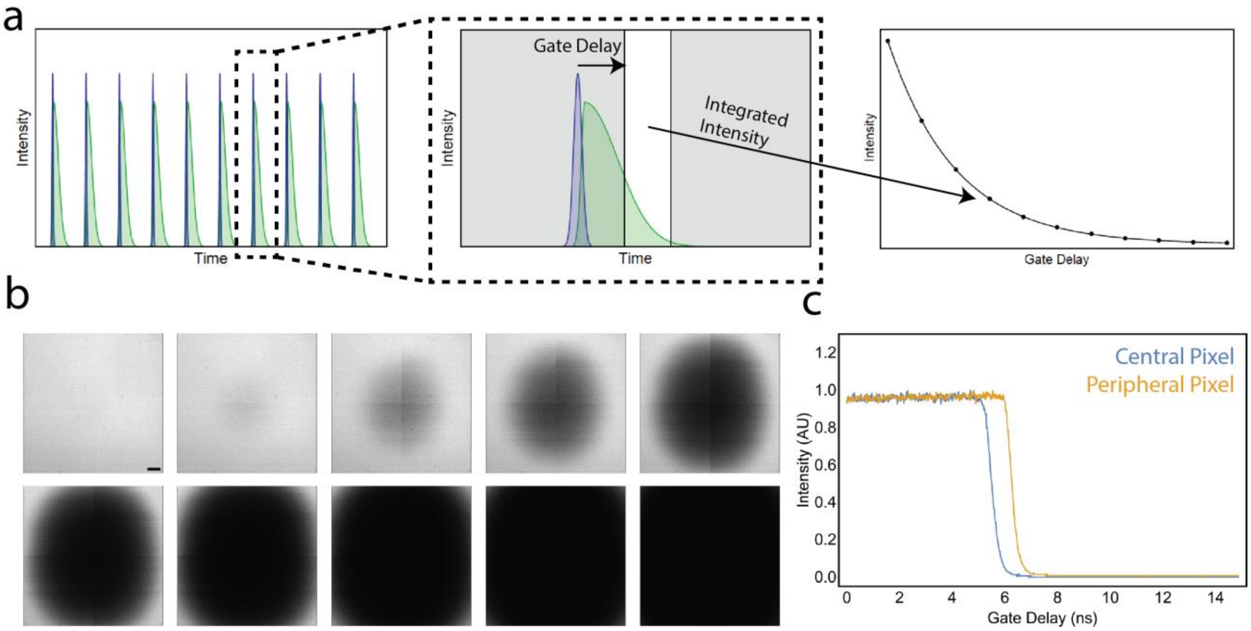


Figure S1: (a) Schematic representation of time-gated imaging. A series of excitation pulses excite a fluorescent molecule, which has a characteristic decay time. The detector collects photons during a “gate” at a specified “gate delay” from the excitation pulse. The signal in this gate is integrated, and the process repeated at increasing gate delays to build up a decay curve. (b) Images from an example instrument response function (IRF) across the full field of view. Time step between images is 18.6 ps, scale bar = 20 µm. (c) Normalized intensity of a single pixel, either central or peripheral, from the IRF plotted over time.

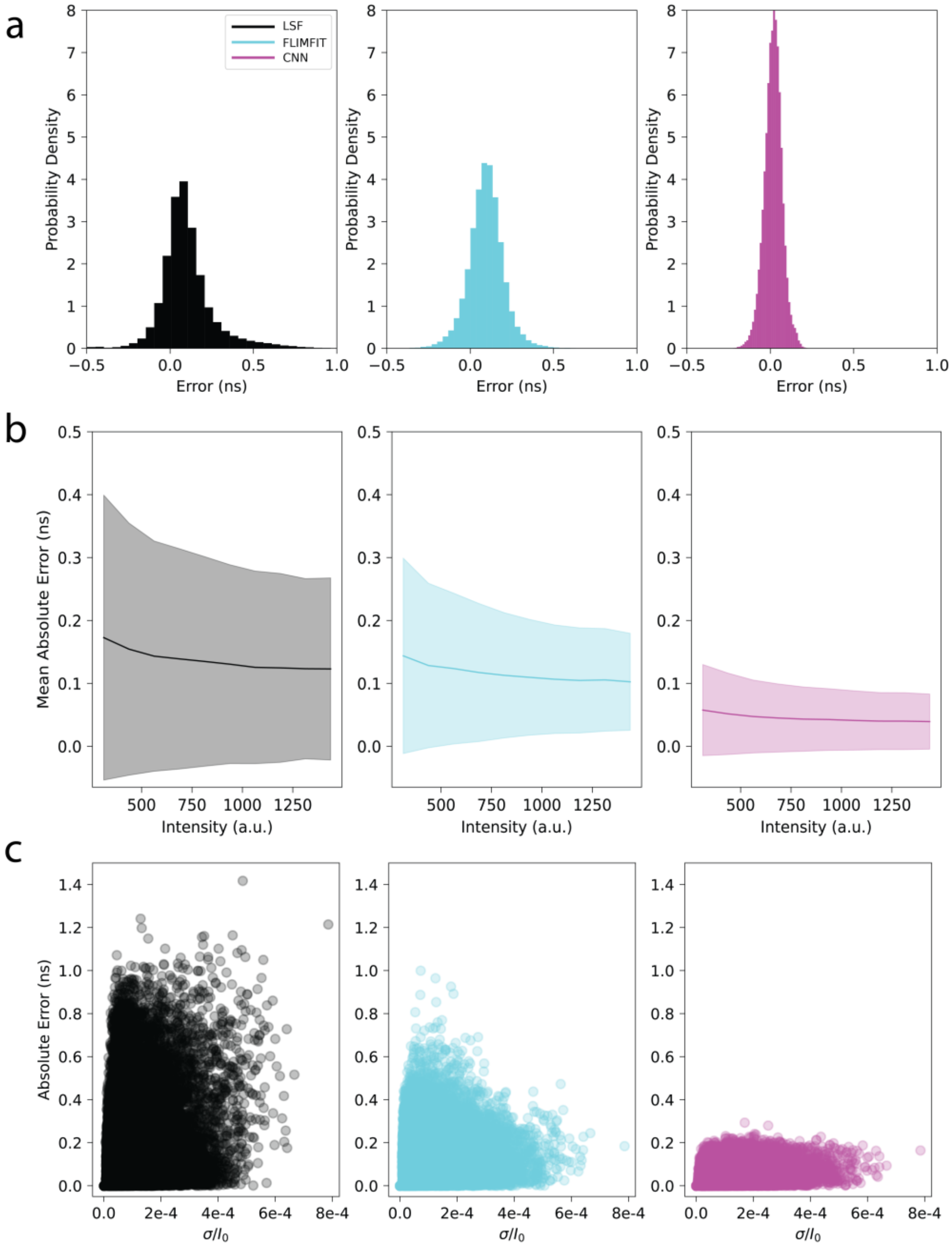


Figure S2: Quantification of lifetime prediction error for simulated decays using three different methods. (a) Absolute error (defined as the absolute value of the difference between the ground truth and prediction) for three methods of lifetime determination: least-squares-fitting (left, black), the python package FLIMFIT (center, cyan), and the ML pipeline (right, magenta). (b) Mean absolute error plotted as a function of intensity for the three methods. Dark line represents the mean error, while the shaded bounds correspond to one standard deviation. (c) Absolute error plotted as a function of the ratio of the standard deviation of the pre-decay signal and the initial intensity. Each point corresponds to a single decay in the test set.

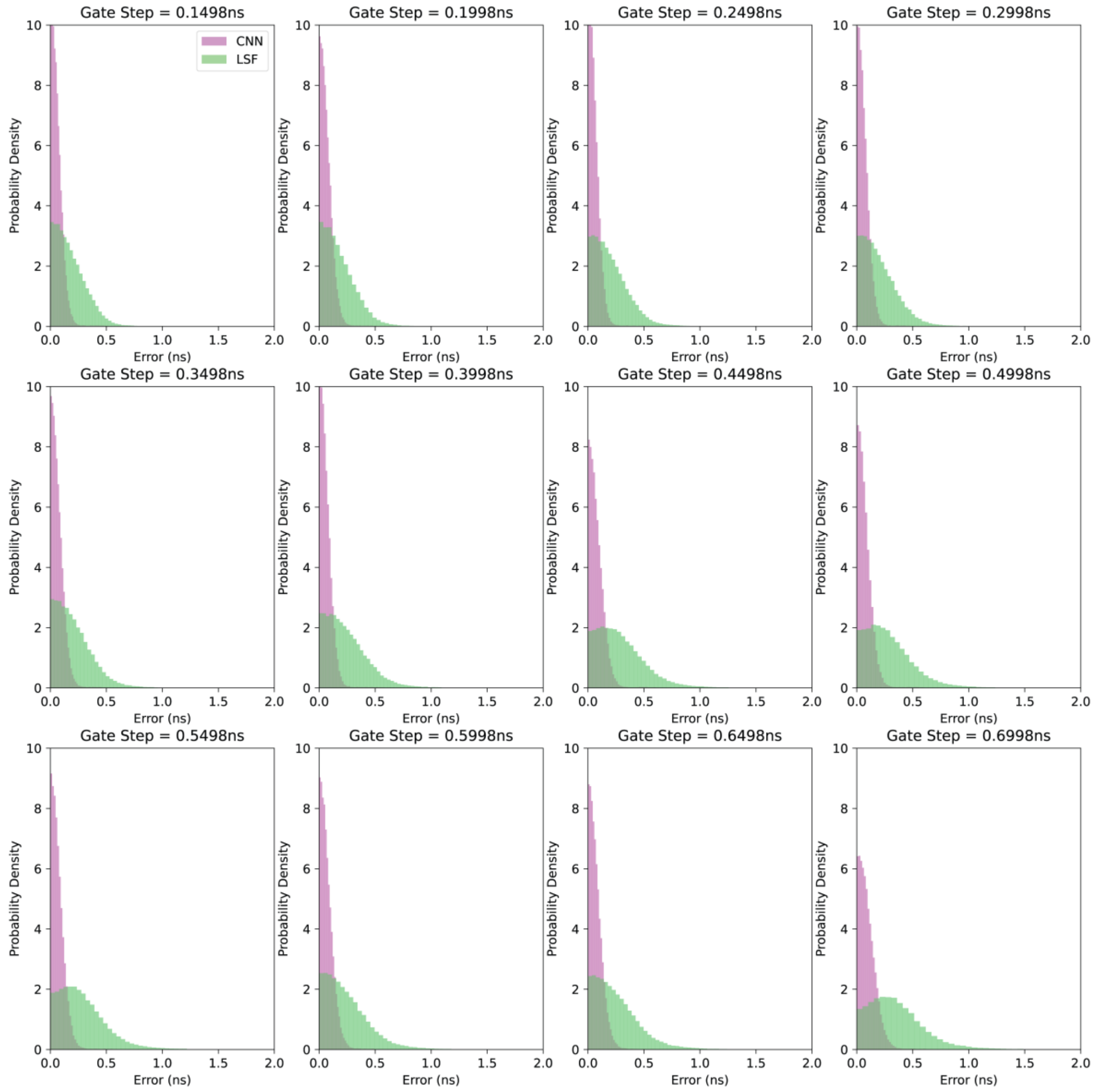


Figure S3: Absolute error plotted as a function of the size of the gate step (which is inversely related to the number of gates) for both the ML pipeline (magenta) and LSF (green). These data reflect the underlying error distributions for the data in Figure 2f.

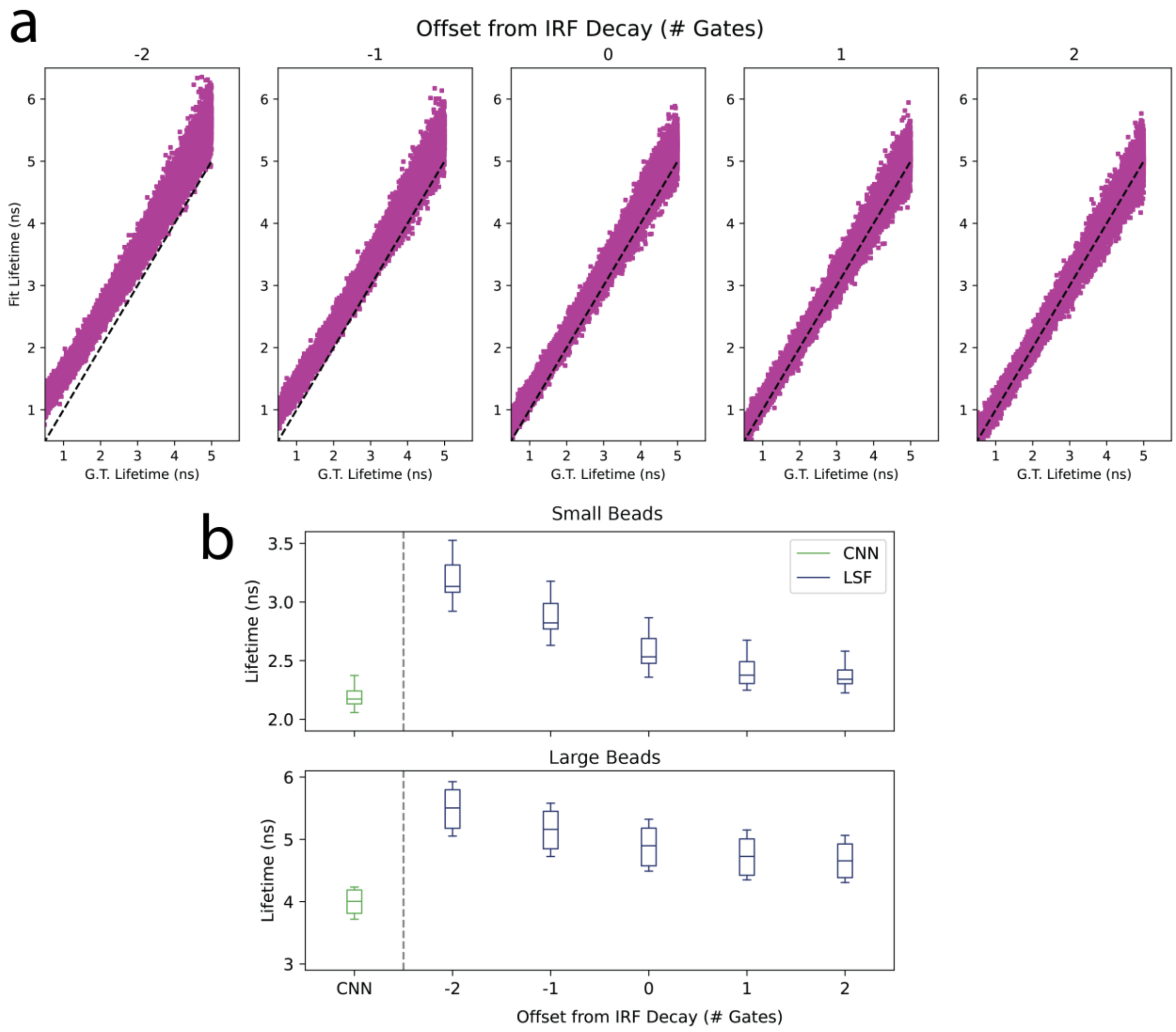


Figure S4: (a) Comparison of the predicted lifetime from least-squares fitting over a range of choices of the start of the decay, given as an integer offset from the calculated start of the IRF decay. (b) Measured lifetimes for the two populations of beads presented in Figure 2a using the ML pipeline and least-squares-fitting. The horizontal line represents the median lifetime. The box spans from the first quartile (Q1) to the third quartile (Q3). The whiskers extend from the box to the farthest data point lying within 1.5x the inter-quartile range (IQR) (adapted from the documentation of matplotlib.pyplot). For the LSF fits, measurements are shown for a range of gates corresponding to the start of the fit, shown as an integer offset from the reference gate in the IRF (see Methods).

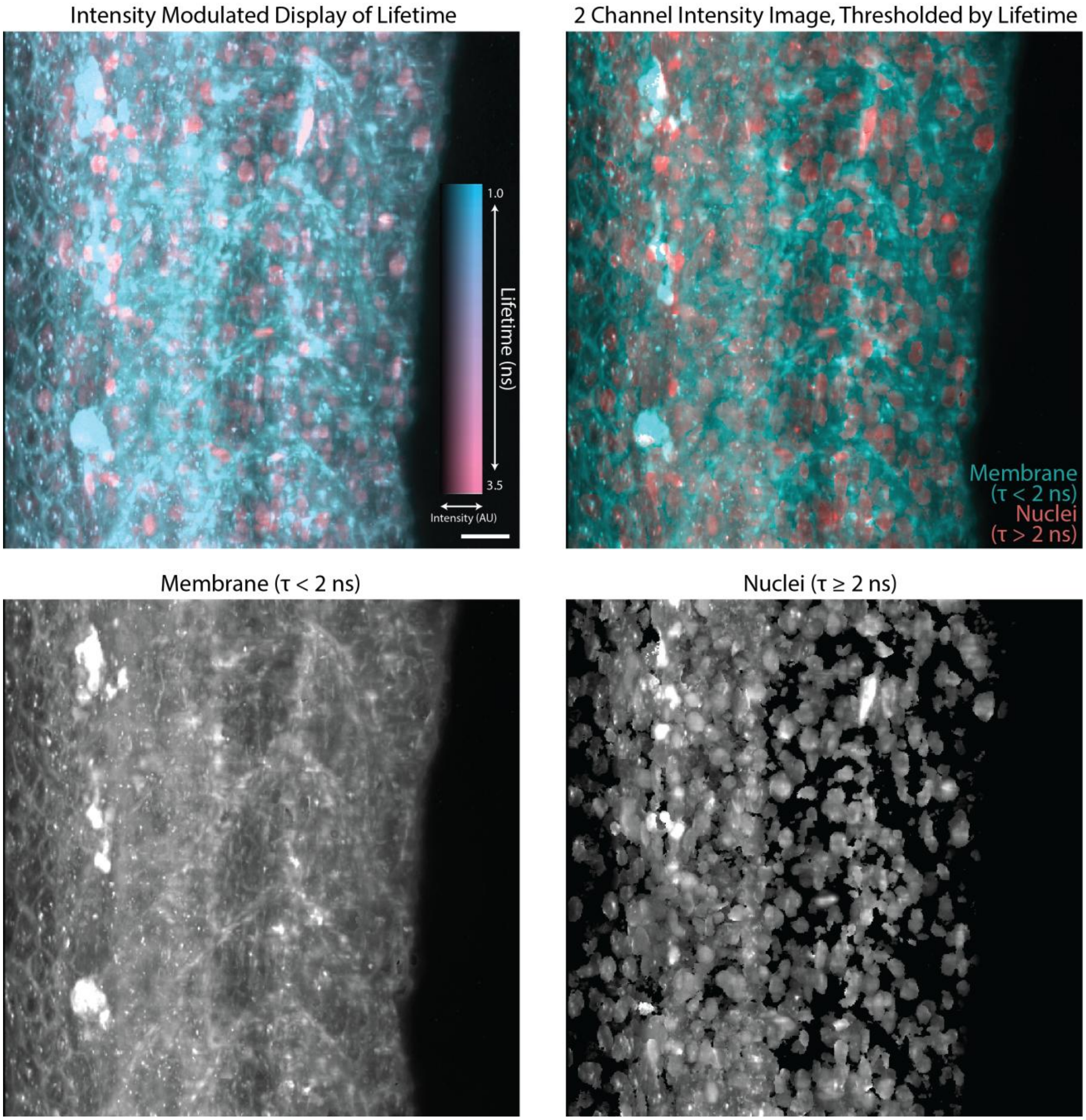


Figure S5: Maximum intensity projections of (top, left) intensity modulated display of lifetimes, (top, right) intensity image thresholded into two channels via lifetime, (bottom, left) intensity image for T < 2 ns, and (bottom, right) intensity image for T ≥ 2 ns. Data set and sample details are the same as in Figure 3d,e. Scale bar = 20 μm.

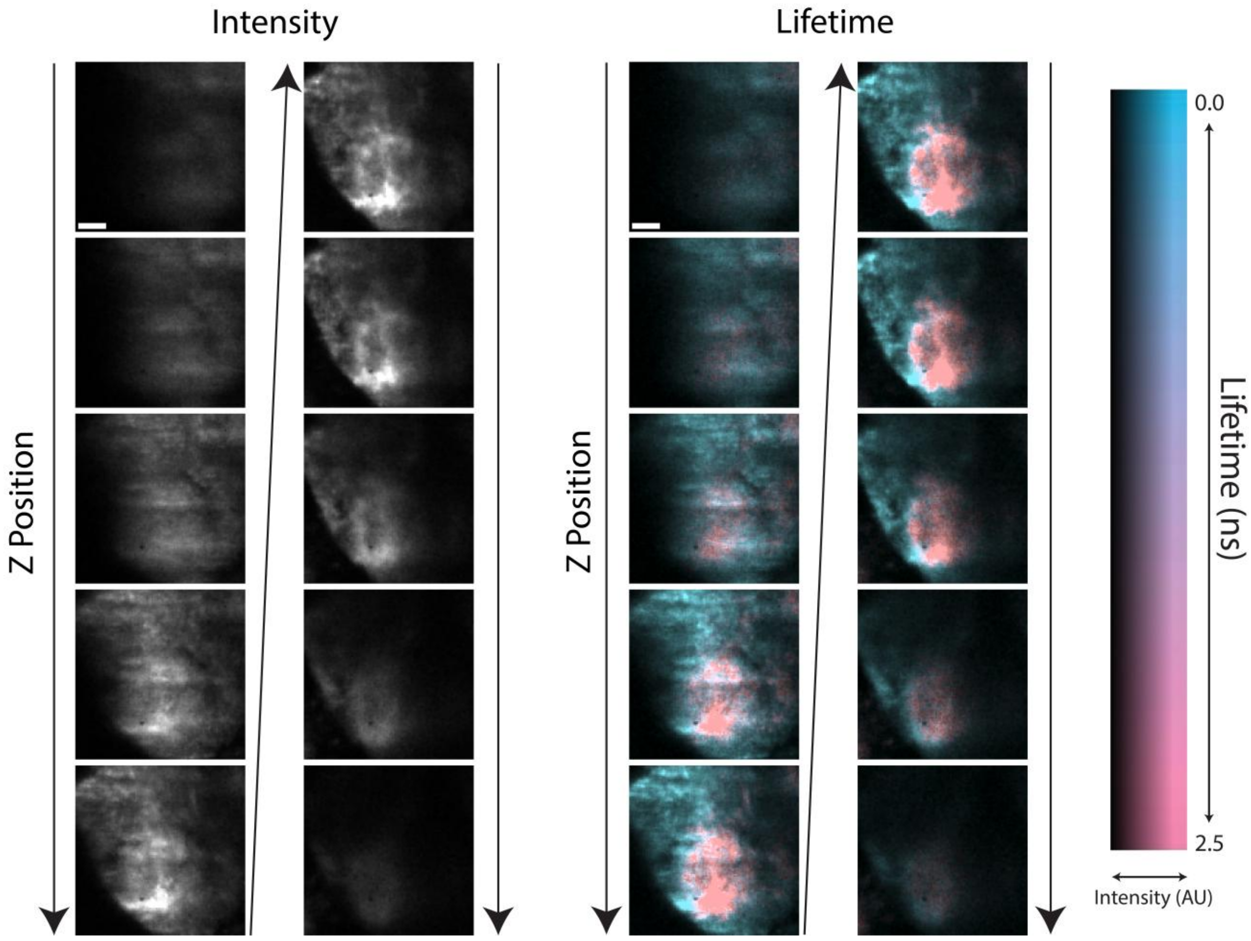


Figure S6: Individual z slices of (left) intensity and (right) intensity modulated display for the area indicated by the white arrow in Figure 3f. Scale bar = 8 μm.

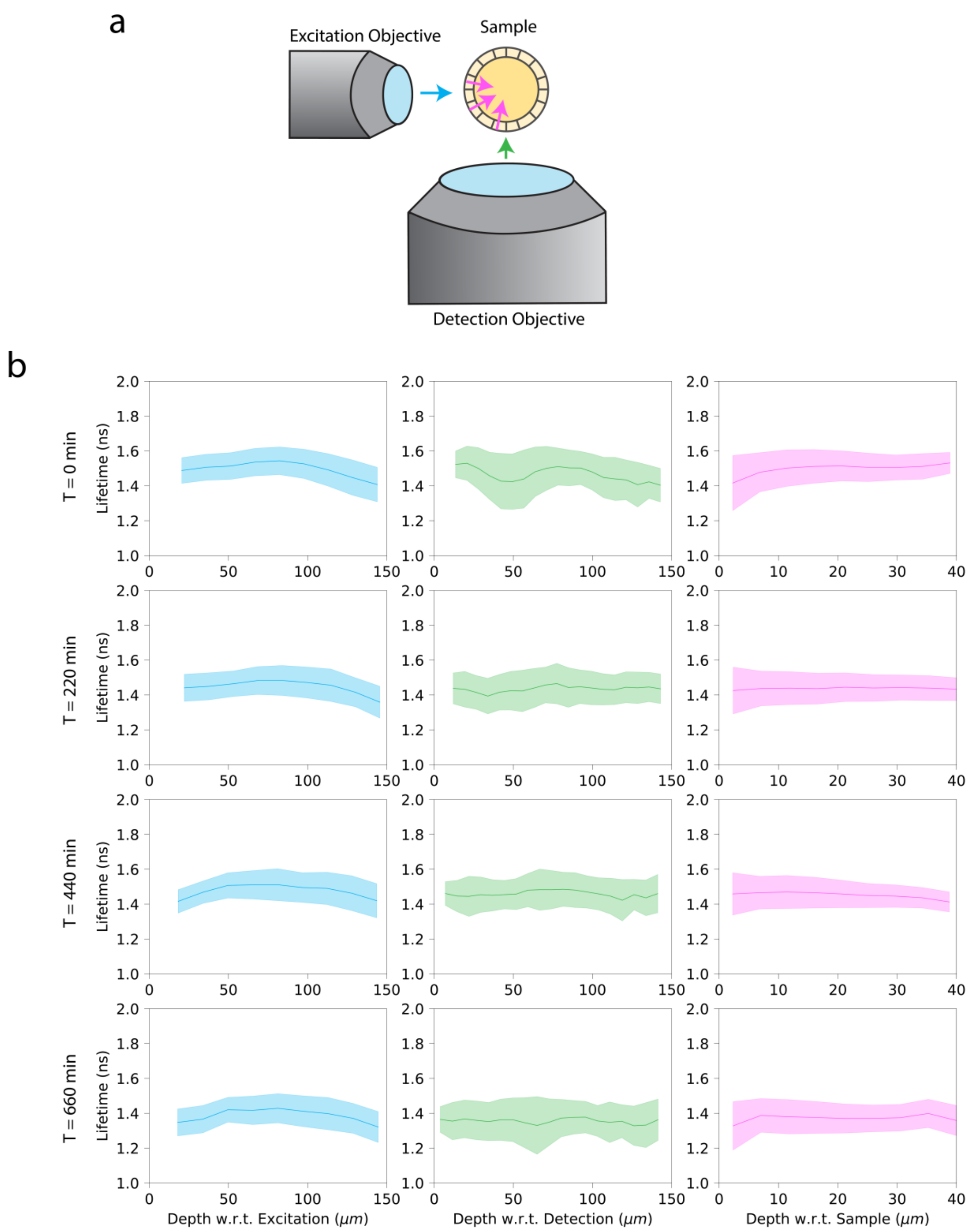


Figure S7: Stability of predicted lifetime as a function of depth in three directions. (a) Schematic indicating three directions of “depth”: from the excitation objective, from the detection objective, and from the surface of the sample itself. (b) Predicted lifetimes plotted as a function of the three depths, at four separate timepoints. The colors correspond to the arrows in (a).

#### Supplemental Movie Captions

Movie S1: 3D rendering of the intensity and intensity modulated display of lifetimes for two populations of fluorescent beads. Data and sample correspond to Figure 3a-c.

Movie S2: 3D rendering and individual z slices of the intensity and intensity modulated display of lifetimes for the trunk of a zebrafish larvae fixed at 4dpf expressing red fluorescent markers for cell membranes and nuclei. Data and sample correspond to Figure 3d,e.

Movie S3: Time series of a 3D rendering of intensity modulated display of lifetimes for the head of a live zebrafish larva at 4 dpf expressing a green fluorescent marker for macrophages. Also detected is tissue autofluorescence. Data and sample correspond to Figure 3f.

Movie S4: Time series of a 3D rendering of intensity modulated display of lifetimes for the head of a live zebrafish larva at 4 dpf expressing a green-red fluorescent photoconvertible marker for macrophages and a red fluorescent marker for blood vessels. Also detected is tissue autofluorescence. Two excitation channels (488 nm and 560 nm) were acquired sequentially per z-stack. Photoconversion was performed at the halfway point of the experiment via an external 375 nm laser line. Data and sample correspond to Figure 4.

Movie S5: Time series of a 3D rendering of intensity modulated display of lifetimes for a *Drosophila melanogaster* embryo developing from approximately stage 6 to stage 16. The embryo expressed a red fluorescent marker for cell nuclei. Data and sample correspond to Figure 5.

Movie S6: Time series of lifetimes for the optic tectum of a life zebrafish larva at 4 dpf expressing a fluorescent reporter for calcium activity. Shown are two individual z slices (scale bar = 10 μm). The average and standard deviation of lifetime for the blue and orange regions are shown over time, highlighting the detection of calcium fluctuations. Data and sample correspond to Figure 6.

### Supplemental Tables

#### Table S1: Sample details, imaging parameters, and processing steps for all experiments.

| Figure | Sample (species, age, region) | Mounting (% agarose, medium) | Fluorescent Labels | Channels | Voxel Size (µm) | Field of view (voxels) | Gate Steps | Gate Step Size (ps) | Gate Width (ns) | Bit Depth | Effective Exposure Time (ms) | Time Points | Temporal Sampling | Destriping? | Visualization |
|---|---|---|---|---|---|---|---|---|---|---|---|---|---|---|---|
| Figure 3a-c; Movie S1 | Fluorescent beads (Bangs Laboratories, #FSDG006, 4.19 µm; Sigma-Aldrich, #L4530-1ML, 2.0 µm) | 2% UltraLowMelting agarose; MilliQ Water | Dragon Green and Unspecified Yellow-Green dye | 488 nm | .4125 x .4125 x 1.0 | 512 x 512 x 211 | 80 | 409 | 20 | 8 Bit | 350 | 1 | N/A | No | Intensity Modulated Display, Imaris |
| Figure 3d,e; Movie S2 | Tg(T2[bactin2::H2B-HaloTag]; myl7:GFP) * Tg(T2[bactin2::mCherry-CAAX]; myl7:GFP); 4dpf; trunk region. HaloTag labelled with $JF_{552}$-HTL | 2% UltraLowMelting agarose; Filtered Fish water | JF552 and mCherry | 560 nm | .4125 x .4125 x 1.0 | 512 x 512 x 101 | 50 | 800 | 20 | 8 Bit | 200 | 1 | N/A | Yes | Intensity Modulated Display, Imaris and FIJI |
| Figure 3f, Movie S3 | Tg(mpeg1:EGFP)gl22; 4dpf; head (hindbrain region) | 2% UltraLowMelting agarose; Filtered Fish water | EGFP | 488 nm | .4125 x .4125 x 2.0 | 512 x 512 x 61 | 30 | 651 | 20 | 8 Bit | 150 | 360 | 30 s | No | Intensity Modulated Display, Imaris |
| Figure 4; Movie S4 | Tg(fli1a:RFP-CAAX)pt505 * Tg(mpeg1:Dendra2)uwm12; 4-5dpf; head (hindbrain region) | 2% UltraLowMelting agarose; Filtered Fish water | RFP1 and Dendra2 | 488 nm; 560 nm | .4125 x .4125 x 2.0 | 512 x 512 x 66 | 30; 30 | 577; 744 | 20; 20 | 8 bit | 150 | 59 | 2 min | No | Intensity Modulated Display, Imaris |
| Figure 5; Movie S5 | w*; P{w[+mC]=His2Av-mRFP}III.1 (Bloomington, 23650), Embryonic stages 7-16 | 2% UltraLowMelting agarose; tap water | mRFP1 | 560 nm | 0.4125 x .4125 x 2.0 | 512 x 512 x 101 per tile, 3 tiles in vertical direction | 30 | 651 | 20 | 8 Bit | 150 | 180 | 4 min | Yes | Intensity Modulated Display, Imaris |
| Figure 6; Movie S6 | Tg(elavl3:NES-WHaloCaMP1a-EGFP) (ZIRC ID: jf820Tg); 4dpf; optic tectum. WHaloCaMP1a labelled with $JF_{552}$-HTL. | 2% UltraLowMelting agarose; Filtered Fish water | JF552 | 560 nm | .4125 x .4125 x 4.0 | 512 x 512 x 5 | 30 | 700 | 20 | 7 Bit | 60 | 115 | 1.04 s | No | Lifetime, FIJI |
